# The Tor pathway, ribosome concentration, and wobble decoding mediate inhibitory effects of the Leu-Pro CUC-CCG codon pair in *Saccharomyces cerevisiae*

**DOI:** 10.64898/2026.02.20.707122

**Authors:** Brandon S. Bruno, Evan M. Platten, Lisa Houston, Christina E. Brule, Elizabeth J. Grayhack

**Author notes:** Corresponding Author: Elizabeth J. Grayhack, Department of Biochemistry & Biophysics, RNA Center, University of Rochester School of Medicine and Dentistry, 601 Elmwood Ave, Rochester, NY 14642, **E-mail**. These authors contributed equally (Brandon S. Bruno, Evan M. Platten). Brandon Bruno Department of Microbiology and Immunology, Stony Brook University Renaissance School of Medicine, 101 Nicolls Road, Life Science Building, Stony Brook NY 11794.

## Abstract

Translation elongation and efficiency are modulated by the genetic code. In the yeast *Saccharomyces cerevisiae,* 17 inhibitory codon pairs, distinguished by requirements for wobble decoding and distinct codon order, result in reduced translation efficiency and slow translation. Nine of these inhibitory pairs are functionally important as they are disproportionately strongly conserved within the orthologous genes in *Saccharomyces sensu stricto*. For three pairs, including CGA-CGA, inhibition is triggered by ribosome collisions and known quality control responses, but the mechanisms by which other pairs cause inhibition is unknown. Here, we examined of the molecular basis of inhibition by the slowly translated, highly conserved Leu Pro CUC-CCG codon pair yielding four findings. First, inhibition is mediated by tRNA^Leu(UAG)^, which decodes CUC by a U●C wobble interaction and effectively competes with the nonessential W●C base pairing tRNA^Leu(GAG)^. Second, despite nearly universal conservation of U33 in tRNAs, the C33 alteration in tRNA^Leu(GAG)^ does not significantly impair its function. Third, inhibition is likely due to ribosome collisions as many suppressors have mutations predicted to reduce ribosome concentration, including mutations in large ribosomal subunit proteins, RNA polymerase I, and ribosome assembly factors. Furthermore, local reduction in ribosome concentration suppresses inhibition. Fourth, we find a link between the metabolic state and CUC-CCG inhibition, as we find six suppressor mutations in *SCH9*, a downstream effector of TORC1 that mediates ribosome production. As Sch9 is inactive during starvation, causing reduced ribosome concentration, one biological function of inhibitory pairs may be to mediate a change in relative expression during starvation conditions.

## INTRODUCTION

Interactions between codons in mRNA and the tRNAs that decode them influence the rate of translation elongation, ultimately controlling both the amount and folding of the nascent polypeptide (Quax et al. 2013; Sander et al. 2014; Tarrant and von der Haar 2014; Pelechano et al. 2015; Yu et al. 2015; Buhr et al. 2016; Brule and Grayhack 2017; Walsh et al. 2020; Liu et al. 2021; Moss et al. 2024). These interactions depend on the particular choice of synonymous codons, which specify insertion of the same amino acid, but are decoded by distinct tRNAs that differ in their abundance as well as in the requirement for wobble interactions at the third base of the codon (Liu et al. 2021; Moss et al. 2024). The use of codons decoded by abundant tRNAs with W●C interactions at all 3 nucleotides (called optimal codons) generally results in efficient expression, while use of suboptimal codons in many organisms, including the yeast *Saccharomyces cerevisiae*, generally results in reduced protein levels and increased mRNA decay (Presnyak et al. 2015; Hanson and Coller 2018; Dowdle and Lykke-Andersen 2025).

The impact of both tRNA abundance and wobble decoding of the A site codon were initially inferred from correlations between codon use, tRNA levels and gene expression (Grantham et al. 1980; Ikemura 1981; Bennetzen and Hall 1982; Ikemura 1982; Curran 1995; dos Reis et al. 2004), yielding models in which competition between cognate and near-cognate tRNAs, or the conflict between supply and demand, mediate codon effects (Elf et al. 2003; Pechmann and Frydman 2013; Dana and Tuller 2014), see (Rodnina 2016). Measurements of codon effects on local translation rates in vivo with ribosome profiling confirmed the importance of tRNA concentration and wobble decoding (Stadler and Fire 2011; Wu et al. 2019); furthermore, models based on measured translation elongation rates confirm that protein output is strongly correlated with predicted translation time (Tunney et al. 2018). However, the effects of codons were inexplicably dependent upon context, with different effects in different locations (Gustafsson 2004; Baranov et al. 2015; Brar 2016).

In fact, codon effects are not only determined by the codon in the A site of the ribosome, but by the identity of the codon in the P site; that is, codon effects are due to combinations of two adjacent codons, called either codon pairs or dicodons. We initially found that, in the yeast *S. cerevisiae*, CGA-CGA codon pairs were much stronger inhibitors of translation than isolated CGA codons (Letzring et al. 2010). This was followed by our identification of 16 other inhibitory codon pairs that confer substantially reduced expression; these pairs are enriched for codons decoded by I●A or U●G wobble interactions, and are in many cases among the most slowly translated codon pairs in the entire native yeast genome (Gamble et al. 2016). Their inhibitory effects almost certainly depend upon the codon pair rather than the component codons, as inhibition by the codon pair is much greater than the sum of the inhibition of the individual codons, and for each of the 12 non-symmetrical inhibitory pairs, the order of the inhibitory codon pairs is crucial to inhibition (Gamble et al. 2016). Inhibitory pairs cause defects in filling the ribosomal A site, based on the observed accumulation of ribosomes with an empty A site at all 17 pairs in vivo, and on in vitro biochemical analysis of elongation rates at two pairs, CGA-CCG and CGA-CGA(Tesina et al. 2020), each of which alters the mRNA conformation to a structure incompatible with decoding (Tesina et al. 2020).

The impact of codon pairs as a regulatory unit is seen also in mutants with defects in the U34 modifications mcm^5^U, ncm^5^U and mcm^5^s^2^U. The crucial role of U34 and other tRNA modifications, inferred from properties of mutants lacking these tRNA modifications (Johansson et al. 2008; Grosjean and Westhof 2016; Phizicky and Hopper 2023; Smith et al. 2024), implicated the U34 modification to mcm^5^s^2^U34 in three tRNAs in both decoding rates at particular codons and protein folding (Nedialkova and Leidel 2015). Subsequent analysis showed that the magnitude by which translation is slowed at some codons decoded by the corresponding hypomodified tRNAs depends upon the identity of the adjacent P site codon, that is the codon pair (Wu et al. 2025). The interplay between codons in the P and A sites was also clearly elucidated in mechanisms of programmed frameshifting (Atkins and Bjork 2009). Thus, the codon pair behaves as a discrete regulatory unit, in which the tRNA-codon interaction in the P site modulates the codon effects in the A site.

The inhibitory codon pairs, like other synonymous codons, are likely biologically important. There is growing evidence for the general importance of synonymous codons, including suboptimal codons, based on observations of conserved rare codon clusters (Chaney et al. 2017; Jacobs and Shakhnovich 2017), and requirements to encode some circadian rhythm regulators with suboptimal codons (Zhou et al. 2013). The importance of the regulation conferred by inhibitory codon pairs is suggested by the extreme conservation of nine of the 17 inhibitory pairs within the *Saccharomyces sensu strictu* yeast. These pairs are much more highly conserved in their locations in orthologous genes than the other codon pairs specifying the same dipeptide and much more highly conserved than expected based on the conservation of their constituent codons (Ghoneim et al. 2019). Thus, we think that inhibitory codon pairs confer a unique and important regulation.

Slow decoding at CGA-CGA codon pairs results in ribosome collisions in vivo, which have been shown to elicit No-Go decay (NGD) (Simms et al. 2017b) with the resulting loss of the mRNA and the protein due to the ribosome quality control complex (Brandman et al. 2012; Letzring et al. 2013). Induction of the NGD pathway by collided ribosomes is conserved from yeast to humans and involves the ubiquitin ligase Hel2 (ZNF598 in humans) (Garzia et al. 2017; Juszkiewicz and Hegde 2017; Matsuo et al. 2017; Sundaramoorthy et al. 2017), the endonuclease Cue2 (NONU-I in *C. elegans*) (D’Orazio et al. 2019; Glover et al. 2020) and the ribosome splitting activity of the RQT complex, including Slh1 and Ascc3 in humans (Matsuo et al. 2017; Juszkiewicz et al. 2020b; Matsuo et al. 2020). Loss of *ASC1*, *HEL2*, *SLH1* or other members of the RQT complex results in increased expression of reporters with CGA codon pairs (Kuroha et al. 2010; Letzring et al. 2013; Veltri et al. 2022). Moreover, the general effects of ribosome collisions as a mechanism of regulation at inhibitory pairs are supported by observations of collided ribosomes at many of these pairs in their native context in vivo (disomes) (Meydan and Guydosh 2020), from documentation that inhibitory codon pairs are associated with increased rates of mRNA decay of native yeast genes (Harigaya and Parker 2017), and from overall low levels of mRNA from these genes in vivo (Chen et al. 2024). However, not all inhibitory codon pairs appear to exert their effects through the same pathways as CGA-CGA, and there are numerous differences between the inhibitory pairs (Table 1). In particular, unlike CGA-CGA, CGA-CGG, and CGA-CCG, which we call the NGD-activating pairs, the inhibitory effects of nine other tested inhibitory pairs are not suppressed by loss of Asc1, and ribosomes do not exhibit frameshifting at these other pairs upon loss of Mbf1 (Wang et al. 2018). Indeed, there are three outstanding questions regarding these codon pairs. What are the properties of tRNAs in A and P sites that cause the inhibitory effects of the pair? What are the mechanisms by which these other inhibitory codon pairs affect translation efficiency? What are the biological functions of inhibitory codon pairs?

**Table 1:** Properties of Inhibitory Codon pairs.

| Codon pair | Di-peptide | Inhibition<br>Median_<br>Syn GFP<sup>1</sup> | Ribo-<br>density<br>rank<sup>1</sup> | Order<sup>1</sup> | Suppression | | Frame-<br>shift in<br>mbf1Δ<sup>2</sup> | Con-<br>served<sup>3</sup> | # Genes with | |
| --- | --- | --- | --- | --- | --- | --- | --- | --- | --- | --- |
| | | | | | tRNA<br>Supp<sup>1</sup> | *asc1* Δ<sup>2</sup> | | | *S. cerevisiae* | Conserved |
| AGG-CGA | RR | 0.48 | 557 | ✓ | ☒ | ☒ | ☒ | ☒ | 129 |  |
| AGG-CGG | RR | 0.82 | 251 | ✓ | nt | nt | nt | ☒ | 101 |  |
| AUA-CGA | IR | 0.58 | 138 | ☒ | nt | nt | nt | ☒ | 268 |  |
| AUA-CGG | IR | 0.65 | 202 | ✓ | nt | nt | nt | ✓ | 164 | 51 |
| CGA-AUA | RI | 0.51 | 3 | ☒ | ✓ | ☒ | ☒ | ✓ | 236 | 129 |
| CGA-CCG | RP | 0.44 | 1 | ✓ | ✓ | ✓ | ✓ | ✓ | 17 | 10 |
| CGA-CGA | RR | 0.44 | 5 | ☒ | ✓ | ✓ | ✓ | ✓ | 20 | 12 |
| CGA-CGG | RR | 0.48 | 6 | ✓ | ✓ | ✓ | ✓ | ✓ | 18 | 10 |
| CGA-CUG | RL | 0.47 | 7 | ☒ | ✓ | ☒ | ☒ | ☒ | 91 |  |
| CGA-GCG | RA | 0.44 | 2 | ✓ | ✓ | ☒ | ☒ | ✓ | 35 | 10 |
| CUC-CCG | LP | 0.44 | 4 | ✓ | ✓ | ☒ | ☒ | ✓ | 61 | 25 |
| CUG-AUA | LI | 0.71 | 449 | ✓ | nt | nt | nt | ☒ | 544 |  |
| CUG-CCG | LP | 0.49 | 19 | ✓ | ✓ | ☒ | ☒ | ✓ | 190 | 47 |
| CUG-CGA | LR | 0.50 | 21 | ☒ | ✓ | ☒ | ☒ | ☒ | 151 |  |
| GUA-CCG | VP | 0.80 | 10 | ✓ | nt | nt | nt | ☒ | 267 |  |
| GUA-CGA | VR | 0.53 | 18 | ✓ | ✓ | ☒ | ☒ | ✓ | 202 | 91 |
| GUG-CGA | VR | 0.60 | 17 | ✓ | ✓ | ☒ | ☒ | ☒ | 75 |  |

**1. (Gamble et al. 2016)**, Median\_syn GFP: Score of(NNN)3-GFP variants with FACS and sequencing; Ribo-density Rank: Ribosome occupancy ranking of 3,721 codon pairs, based on 100 codon window in yeast genes; Order: Arrangement of two codons is or is not required for inhibition; tRNA Suppression: Overexpression of 5' tRNA or expression of non-native tRNA restores expression of GFP variant with the inhibitory pair
**2. (Wang et al. 2018)** *asc1* $\Delta$ : *ASC1* deletion results in increased expression of reporters with inhibitory pair relative to that of reporters with the optimal pair; Frameshift in *mbf1* $\Delta$ : *MBF1* deletion results in +1 frameshifting at the codon pair
**3. (Ghoneim et al. 2019)** Conserved: Conservation of the inhibitory codon pair at the same location within paralogous genes across 5 *Saccharomyces sensu stricto* species is more than 3 standard deviations greater than expected based on conservation of their individual codons; Number of genes with each individual inhibitory pair in *S. cerevisiae* (L) or conserved location across 4 of the 5 *sensu stricto* species.

To address these questions, we studied the strongly inhibitory (Leu-Pro) CUC-CCG codon pair. While the contribution of U●G wobble decoding of CCG was previously demonstrated (Gamble et al. 2016), the question of what aspect of CUC decoding is responsible for its inhibitory effects in this codon pair is particularly intriguing. Two tRNAs are implicated in decoding CUC: tRNA^Leu(GAG)^, with an anticodon that is an exact base pairing match for the CUC codon and with C33, instead of the nearly universally conserved U33 in the anticodon loop, is encoded by a single copy nonessential gene (*tL(GAG)*; and tRNA^Leu(UAG)^, which requires an U●C wobble interaction to decode CUC is encoded by three genes (*tL(UAG)* (Farabaugh et al. 2006; Johansson et al. 2008; Grosjean and Westhof 2016). We provide evidence by manipulating tRNA genes that both tRNAs decode CUC effectively, but that tRNA^Leu(UAG)^ confers strong inhibition in CUC-CCG pairs, implicating wobble interactions in the P site as causative for inhibition.

To understand the mechanism(s) by which CUC-CCG codons cause reduced protein expression, we selected suppressors of CUC-CCG inhibition and identified the responsible mutations by whole genome sequencing, and appropriate reconstruction experiments. In this way, we found that CUC-CCG inhibition is suppressed by mutations in genes expected to affect ribosome abundance (based on mutations in four *RPL* genes encoding one of two duplicated copies of large subunit ribosomal proteins; *RPA190*, encoding the largest subunit of RNA polymerase I; and four genes encoding proteins involved in ribosome biogenesis). We then provide additional evidence that reduction in translating ribosomes on a particular transcript suffices to suppress inhibition. Furthermore, we provide evidence that the reduction in large subunit proteins results in reduced inhibition by at least 6 inhibitory pairs, implying that it is a general mechanism. These results suggest CUC-CCG inhibition does operate through ribosome collisions. As we also obtained 6 mutations in the TORC1-regulated kinase *SCH9*, which regulates both ribosomal protein and rRNA expression, we propose that the effects of this and perhaps other inhibitory codon pairs depend upon nutritional status of the cell and thus, that inhibitory codon pairs may confer a coherent regulation on genes in which they occur.

## Results

### A reporter system in which each CUC-CCG codon pair exerts similar levels of inhibition

We chose to study inhibition by the CUC-CCG codon pair, encoding Leu-Pro, one of the nine pairs that differ from CGA-CGA, based on four properties of this pair (see Table 1). First, CUC-CCG codon pairs are strongly inhibitory, comparable to the CGA-CGA codon pair (encoding Arg-Arg), based on both high throughput scoring of multiple GFP variants and on assessment of individual reconstructed variants (Gamble et al. 2016). Second, CUC-CCG is slowly translated in yeast based on ribosome density (the 4^th^ most slowly translated pair, equivalent to CGA-CGA) and enrichment in disomes (Meydan and Guydosh 2020), indicating that ribosome elongation is fundamentally compromised at this pair. Third, CUC-CCG is likely biologically important as the codon pair is one of nine highly conserved inhibitory pairs within the *Saccharomyces sensu strictu* species (present in the same genes at the same locations in five species) (Ghoneim et al. 2019). Fourth, CUC-CCG is one of only three highly conserved codon pairs that lack a CGA codon and the basis for CUC inhibitory effects are unknown (Farabaugh et al. 2006; Johansson et al. 2008; Grosjean and Westhof 2016).

To study inhibition by the CUC-CCG inhibitory pair, encoding the Leu-Pro dipeptide, we designed a reporter to recapitulate natural inhibitory conditions and still allow us to amplify the signal with multiple copies of an inhibitory pair. Most genes contain only a single inhibitory pair (Ghoneim et al. 2019), although inhibitory codon pairs are relatively common, found in ∼30% of yeast genes (Gamble et al. 2016). Thus, to study multiple independent events that each might require ribosome collisions, we designed a reporter in which the codon pairs were inserted far enough into the coding sequence to allow efficient ribosome collisions (Simms et al. 2017b) and were separated from each other so that each inhibitory event was independent of preceding events (collisions) (Fig. 1A). To this end, we fused two non-identical copies of the 10th fibronectin type III domain of human fibronectin (FN-FN) to GFP, with the first codon pair inserted at amino acid 111 and subsequent pairs separated by at least 31 codons (Fig. 1A). We chose the10th fibronectin type III domain of human fibronectin (FN), which is composed of 7 β sheets, because there is extensive documentation that insertions in its loop regions generally do not affect the β sheet structure (Koide and Koide 2007) and because it is unlikely that yeast will exert selection on the human sequence. The arrangement of inhibitory and optimal codons are designated by the position of the insert, thus OOOO specifies a reporter in which all four insertions encoding Leu-Pro are optimal codon pairs (UUG-CCA), while IOIO specifies a reporter in which positions 1 and 3 are inhibitory codon pairs (CUC-CCG) and positions 2 and 4 are optimal codon pairs (UUG-CCA).

**Figure 1.**
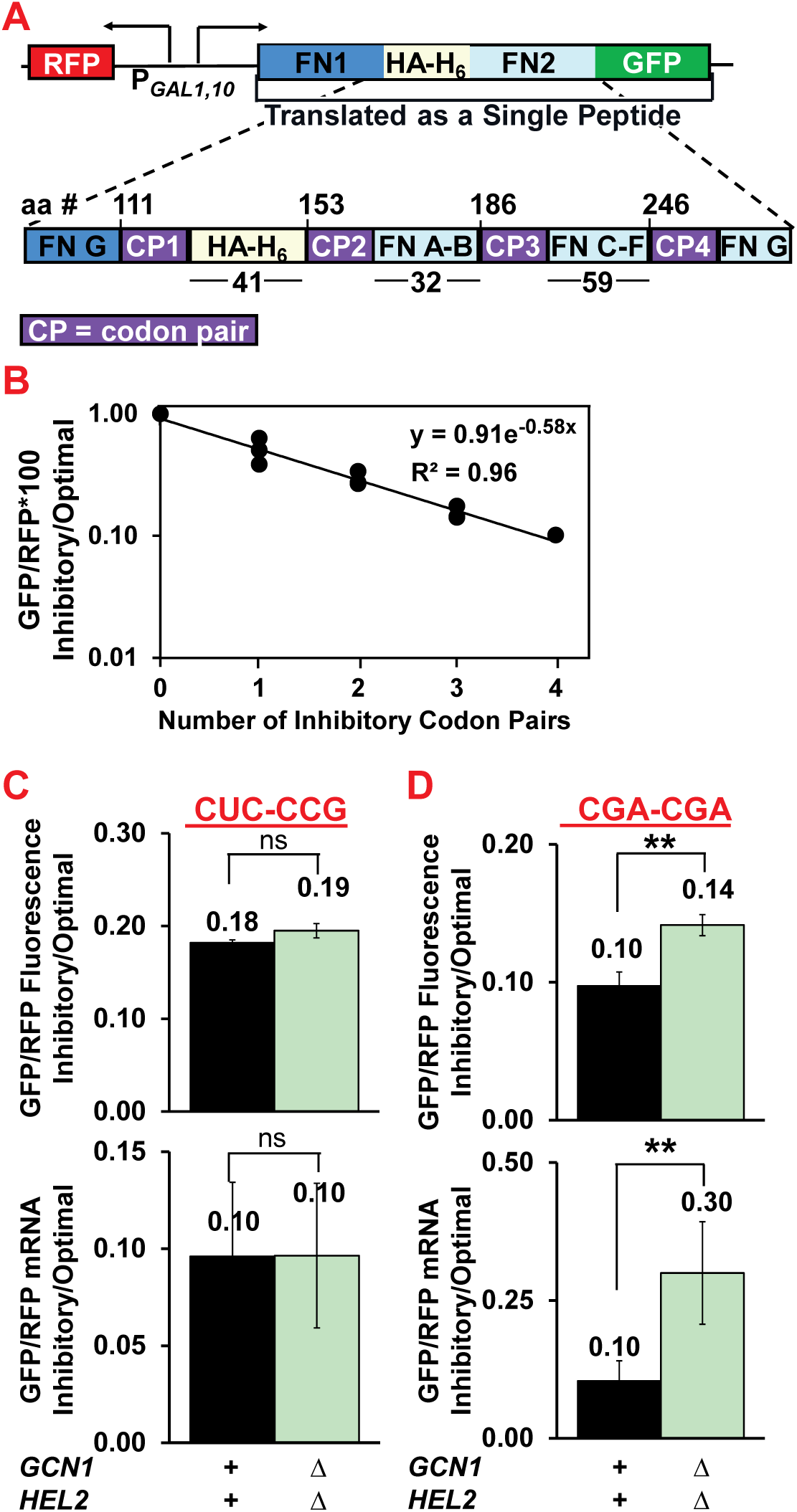
Inhibition by the CUC-CCG codon pair is dose dependent and not responsive to mutations in *HEL2* and *GCN1*. **A.** Diagram of the RFP (red) and FN1 (dark blue)-HA-His_6_ (yellow)-FN2 (light blue)-GFP (green) reporter with expression of both RFP and GFP driven by the bidirectional *GAL1,10* promoter. The FN1-HA-His6-FN2-GFP encodes a single contiguous polypeptide, with the two FN domains differing in their nucleotide sequence to avoid recombination. The expansion below illustrates sites for insertion of codon pairs (CP1-CP4) with the first insertion at the end of the FN1 (tenth fibronectin FNIII) domain (at amino acid 111), followed by insertion downstream of an HA_epitope_-His_6_ sequence (amino acid 153), then into B-C (amino acid 186) and F-G (amino acid 246) loops of the second fibronectin domain (FN2). The insertion sites for inhibitory codon pairs (purple boxes) are separated by the indicated number of codons, shown below the diagram. The sequence of the parent vector EEVN391 is reported in Supplemental Figure S17. **B.** Inhibition by CUC-CCG exhibits exponential dose-dependence independent of position of inhibitory pairs. Each point represents data from a distinct construct, with the graph generated from yeast with one construct with all optimal pairs (OOOO), three different constructs bearing a single inhibitory pair (IOOO; OOIO, OOOI, four constructs with two inhibitory pairs (IOOI, IOIO, OIIO, OOII), three constructs with three inhibitory pairs (IOII, IIIO, OIII) and one construct with four inhibitory pairs (IIII). Optimal pairs are used at all other sites. For each construct, GFP/RFP measurements were the average of three or four independent yeast transformants bearing chromosomally integrated copies of the particular reporter. **C.** CUC-CCG dependent reduction in protein and mRNA is not suppressed in a *hel2Δ gcn1Δ* mutant. Optimal constructs (OOOO) contain four copies of UUG-CCA in FN reporter, while inhibitory constructs have three copies of the inhibitory Leu-Pro CUC-CCG and one copy of UUG-CCA in the FN reporter (IOII arrangement). **D.** CGA-CGA dependent reduction in both protein and mRNA is suppressed in a *hel2Δ gcn1Δ* mutant. Optimal constructs bear three copies of Arg-Arg CGU-CGU and one copy of Leu-Pro UUG-CCA (position 2) in FN reporter (OO_LP_OO), while inhibitory constructs bear three copies of CGA-CGA and one copy of UUG-CCA in the FN reporter (IO_LP_II), as we made the unexpected discovery that insertion of RR at both positions 2 and 3 impaired expression regardless of whether or not the RR was encoded by optimal or suboptimal codons. ** P value <0.01 ≥0.001.

Using this reporter, we found an exponential relationship between inhibition and the number of CUC-CCG inhibitory codon pairs (Fig. 1B), which is the expected result for independent inhibitory events acting in series; we had previously observed this exponential relationship with CGA-CGA codon pairs (Letzring et al. 2010). As described previously, we measure expression based on relative amounts of GFP to RFP as the bi-directional *GAL1,10* promoter directs expression of both proteins (Fig. 1A). We report the expression of a reporter with inhibitory pairs relative to expression of a synonymous reporter with all optimal pairs to facilitate comparisons between different genetic backgrounds. We note that here, as in previous results, the exponential coefficient is indicative of how much each pair reduces expression. As seen in Fig. 1B, we demonstrated that inhibition was independent of the positions of the inhibitory pairs, as we measured GFP/RFP from at least three different constructs with the same number of inhibitory pairs (1, 2 or 3) and found little difference between constructs with the same number of inhibitory pairs. For instance, of our reporters with two inhibitory pairs, three (IOOI; OOII; OIIO) yielded identical GFP/RFP inhibitory/optimal (0.27) and the fourth (IOIO) yielded 0.34; and we obtained an R^2^ value for the fit of 0.96 for the overall fit. This data provides strong evidence that the effect of the inhibitory pair at each position is nearly identical.

### CUC-CCG inhibition is not suppressed by removal of both Hel2 and Gcn1, each of which recognizes collided ribosomes

Recognition of collided ribosomes in yeast is mediated independently by both Hel2 and Gcn1 proteins, which compete with each other with respect to activation of the Integrated Stress Response (ISR) pathway (Meydan and Guydosh 2020; Yan and Zaher 2021; Kim and Zaher 2022), but act cooperatively to prevent frameshifting at CGA codons (Houston et al. 2022). Thus, we considered that inhibition of GFP expression by CUC-CCG could be mediated through both pathways, with either sufficing for complete inhibition. To compare effects of the *gcn1Δ hel2Δ* mutant to wild type on CUC-CCG inhibition, we examined GFP/RFP from both the inhibitory and optimal constructs in both strains, and report the ratio of inhibitory to optimal in the main figure and the individual GFP/RFP values in supplemental figures.

The *gcn1Δ hel2Δ* strain did not suppress inhibition by CUC-CCG codon pairs, as the ratio of GFP/RFP protein (fluorescence) and mRNA levels of inhibitory/optimal reporters is nearly identical in the wild type and *gcn1Δ hel2Δ* strains (Fig. 1C, Supplemental Fig. S1). By contrast, the *gcn1Δ hel2Δ* strain did suppress inhibition by the CGA-CGA codon pairs in the same FN reporter construct (Fig. 1D; Supplemental Fig. S2). Thus, while inactivation of the NGD pathway and the ISR pathway suppresses inhibitory effects of separated CGA-CGA inhibitory pairs, inactivation of these pathways does not suppress inhibition by CUC-CCG codon pairs.

### Inhibition by CUC-CCG is mediated by a competition between two tRNAs that decode CUC *in vivo* and by U●C wobble interactions

To understand why the CUC-CCG codon pair is inhibitory, we needed to determine the identity of tRNAs that decode each codon and the salient properties of these tRNAs. We already knew that U●G wobble decoding of CCG was key to inhibition as expression of a tRNA^Pro(UGG→CGG)^ variant (*tP(UGG) U34C*) (that allows W●C base pairing at all three bases of the codon) suppressed inhibition by CUC-CCG (Gamble et al. 2016). We were particularly interested in defining the relevant interactions for the CUC codon, as either of two tRNAs can decode CUC in the yeast *S. cerevisiae* (Farabaugh et al. 2006; Johansson et al. 2008; Grosjean and Westhof 2016). *S. cerevisiae* encodes a single copy tRNA^Leu(GAG)^ (that can decode CUC by W●C base pairing at all three positions), but this tRNA is not essential and bears a C at position 33 in the anticodon loop, a position that is almost universally a U. While tRNA^Leu(GAG)^ is expressed and its anticodon is functionally important (as its loss suppresses a deletion of *dcp2*) (Kim and van Hoof 2020), it is unknown if C33 impairs tRNA function, as U33 is involved in a U turn important for the conformation of the anticodon loop (Ashraf et al. 1999); moreover, G33 in a *C. albicans* tRNA^Ser^ alters the structure of the anticodon arm and reduces its decoding efficiency, which was arguably key to reassignment of CUG from leucine to serine in a fungal CTG clade that includes *C. albicans* (Santos et al. 1996; Perreau et al. 1999). In addition, *S. cerevisiae* encodes three copies of tRNA^Leu(UAG)^, which is presumed to decode CUC in the absence of tRNA^Leu(GAG)^, but tRNA^Leu(UAG)^ decoding of CUC requires an unusual U●C wobble base pair. Thus, it is unknown if the CUC codon is primarily decoded by tRNA^Leu(GAG)^ and if so, whether tRNA^Leu(GAG)^ has an unfavorable interaction in the ribosome due to C33, or by tRNALeu(UAG) via an unusual U●C wobble.

To determine how the inhibitory properties of CUC-CCG were affected by tRNA^Leu(GAG)^, tRNA^Leu(UAG)^ and the C at position 33 in tRNA^Leu(GAG)^, we examined dose-dependent inhibition by CUC-CCG in strains bearing deletions, insertions or mutations in these tRNA genes (Fig. 2A, B, C). In each case, we compared otherwise isogenic strains, determined the exponential coefficient for inhibition and converted that into the expected inhibitory effect of a single CUC-CCG codon pair (Fig. 2F). We also directly compared the impact of *tL(GAG)* copy number and *tL(GAG)C33U* variants by examining expression of two reporters (containing either two or four inhibitory pairs) in strains with zero, one or two copies of *tL(GAG)* at the native and *LYS2* loci, with C33 and U33 as indicated (Fig. 2D, E). We draw four conclusions from this analysis.

**Figure 2.**
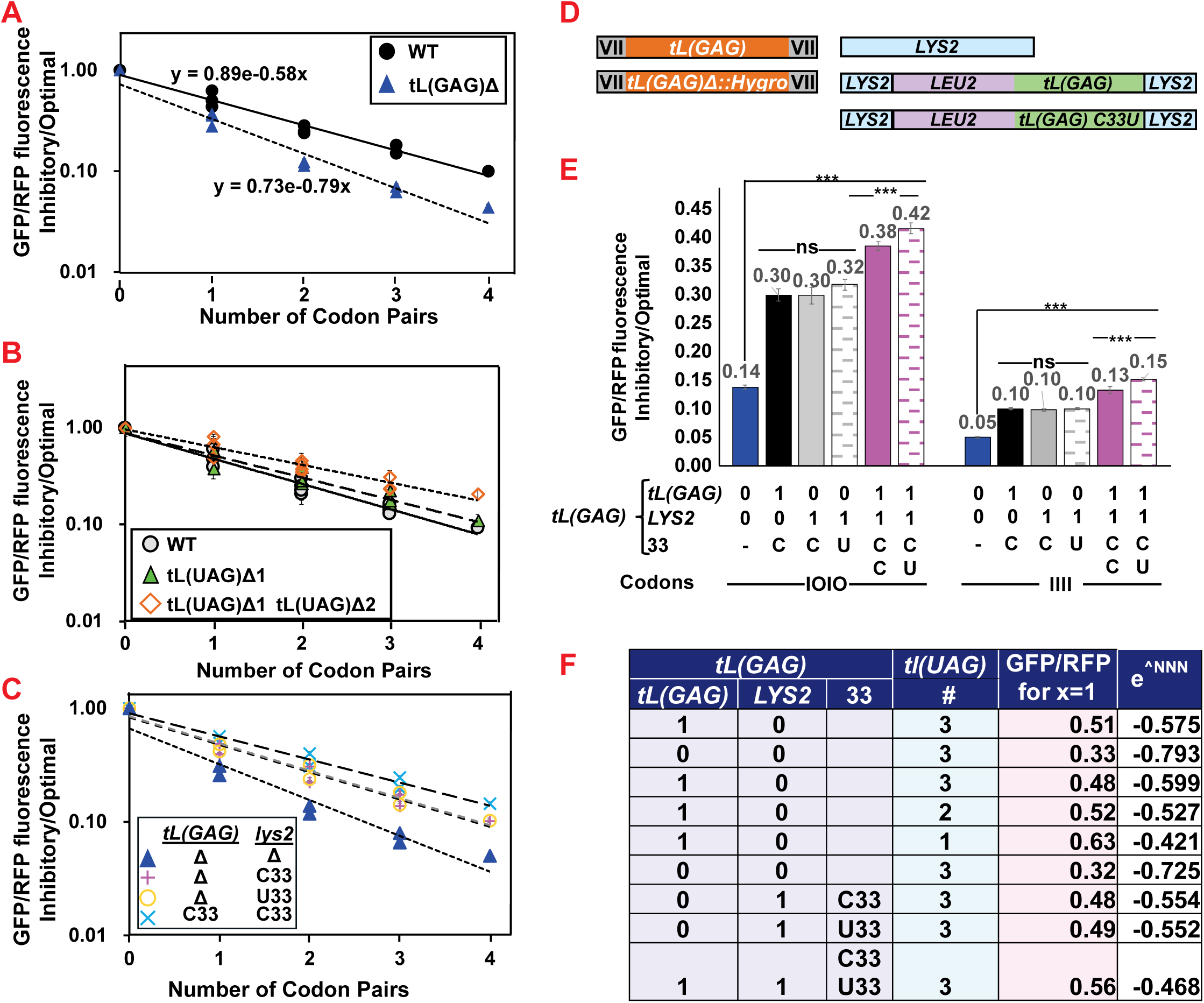
Inhibitory effects of the CUC-CCG codon pair depend upon competition between tRNA^Leu(GAG)^ and tRNA^Leu(UAG)^. **A.** Deletion of the single copy tRNA *tL(GAG)* results in increased inhibition by CUC-CCG. Dose dependent inhibition of FN-GFP/RFP was measured in a wild type strain and an otherwise isogenic *tL(GAG)Δ* strain transformed with FN reporters bearing zero to four copies of the inhibitory CUC-CCG codon pair with the optimal codon pair UUG-CCA at the remaining sites. In addition to OOOO and IIII constructs, data was generated from multiple constructs with one, two and three inhibitory pairs (one pair: OOIO, IOOO, OOOI), (two pairs: OOII, I00I, OIIO), (three pairs: IOII, IIIO). **B.** Deletion of one or two copies of the three copy tRNA *tL(UAG)* results in reduced inhibition by CUC-CCG. Dose dependent inhibition of FN-GFP/RFP was measured in strains bearing deletions of one of the three copies *tL(UAG)* (*tL(UAG)ΔJ)* or two of three copies of *tL(UAG)* (*tL(UAG)ΔJ* (*tL(UAG)ΔL1)*. The parent strains bearing a plasmid borne copy of *tL(UAG)J* were transformed with FN reporters with zero to four copies of CUC-CCG (as above), streaked on FOA to select for loss of the plasmid with *tL(UAG)J*, followed by growth in galactose and analysis of GFP/RFP. **C.** A second copy of *tL(GAG)C33* or *tL(GAG)U33* results in reduced CUC-CCG inhibition. Dose dependent inhibition of FN-GFP/RFP was measured in strains with zero, one or two copies of *tL(GAG)* wt (C33) or one copy of mutant *tL(GAG)*-C33U, with *tL(GAG)* and variants inserted as *LYS2* (see D.) **D.** Diagram of *tL(GAG)* wild type deletion constructs at its native chromosomal location and of insertions of *tL(GAG)* wild type (C33) and the mutant form C33U at *LYS2*. **E.** Comparison of expression of two reporters with two (IOIO) or four (IIII) inhibitory CUC-CCG codons in strains with zero, one or two copies of the tRNA^Leu(GAG)^ at different chromosomal locations and with either C33 or U33. P values, calculated as described previously (Wang et al. 2018) were used to assess significance of differences between all strains and the strain lacking tL(GAG), and to assess differences between strains with the same copy number of tL(GAG). *** P value <0.001. **F.** Comparison of inhibitory effects of CUC-CCG codon pairs in strains with different copy numbers of tRNA^Leu(GAG)^ or tRNA^Leu(UAG)^. The table shows the inhibitory effect of a single CUC-CCG codon pair (calculated based on the exponential and fit to dose dependent curve) in strains bearing different numbers and variants of *tL(GAG)* and *tL(UAG)*. For each strain the table delineates the number, location and N33 status of *tL(GAG)* genes and the number of tl(UAG) genes,

First, we conclude that tRNA^Leu(GAG)^ does decode CUC in vivo and is not primarily responsible for inhibitory effects of the pair, as we find that CUC-CCG inhibition is significantly stronger when *tL(GAG)* is deleted (Fig. 2A, F). The dose curve yields an exponential coefficient of -0.58 in a wild type strain compared to -0.79 in a strain lacking *tL(GAG)* (Fig. 2A). Based on these exponential coefficients, we estimate that a single CUC-CCG reduces expression to 0.51 in the wild type strain, while a single CUC-CCG codon pair reduces expression more strongly to 0.33 in the *tL(GAG)*Δ strain (Fig 2A, 2F). We have performed four repeats of this entire comparison, yielding a compiled estimate that a CUC-CCG pair reduces expression to 0.48 ±0.02 per CUC-CCG pair in the wild type and to 0.32±0.01 in the *tL(GAG)*Δ (p value < E-05). As expected, we found that the CUC-CCG codon pair resulted in concomitant loss of protein and mRNA in strains lacking *tL(GAG)* (Supplemental Fig. S3). These results provide direct evidence that tRNA^Leu(GAG)^ decodes a substantial fraction of the CUC codons in the CUC-CCG codon pair and is not primarily responsible for inhibitory effects of the pair.

Second, we conclude that tRNA^Leu(UAG)^ does decodes CUC (even in the presence of tRNA^Leu(GAG)^) and plays a major role in inhibition, as we find that CUC-CCG inhibition is alleviated by reduction in tRNA^Leu(UAG)^ concentration in a wild type strain (Fig. 2B, 2F). Based on the exponential coefficients, strains lacking either one or two of the three copies of *tL(UAG)* exhibit reduced inhibitory effects of a single CUC-CCG codon pair from 0.48 in the wild type strain to 0.52 and 0.63 respectively in strains lacking one and two of the genes encoding tRNA^Leu(UAG)^. We note, however, that even at an extrapolated value of zero copies of *tL(UAG)*, the CUC-CCG codon pair is still predicted to inhibit expression (to ∼0.7 for a single pair), and thus we cannot state that tRNA^Leu(UAG)^ is solely responsible for inhibition by CUC-CCG (Supplemental Fig. S4). These findings imply that the U●C wobble does not substantially interfere with acceptance of tRNA^Leu(UAG)^ into the A site, as this tRNA must decode a substantial fraction of the CUC codons to account for its effects on inhibition by CUC.

Third, we infer that it is the effective competition between tRNA^Leu(GAG)^ and tRNA^Leu(UAG)^ to occupy the A site that is largely, but not solely, responsible for inhibitory effects of CUC-CCG, as we find that CUC-CCG inhibition is alleviated by increased copy number of *tL(GAG)* (Fig. 2C). Based on the exponential coefficients derived from dose curves in Fig. 2C and shown in 2F, we infer that while a single CUC-CCG codon pair reduces expression to 0.32 in the *tL(GAG)Δ* strain, a single CUC-CCG is less inhibitory in strains with one or two copies of *tL(GAG)*, reducing expression to 0.48 and 0.56 respectively (Fig.2F). Similarly, we observe increased expression of reporters with two or four CUC-CCG pairs with increased copy number of *tL(GAG)*, finding an approximately linear relationship between copy number and expression (Fig. 2E; Supplemental Fig. S5). Thus, we infer that there is a competition between tRNA^Leu(GAG)^ and tRNA^Leu(UAG)^ to occupy the A site, and that occupancy by tRNA^Leu(UAG)^ results in stronger inhibitory effects of CUC-CCG. However, as increased copy number of *tL(GAG)* does not fully suppress the inhibitory effects of CUC-CCG codon pairs, we infer that even with two copies of *tL(GAG)* either tRNA^Leu(UAG)^ still decodes a significant fraction of the CUC codons or tRNA^Leu(GAG)^ is also inhibitory.

Fourth, we infer that C33 has a minimal effect on either tRNA^Leu(GAG)^ function or concentration. Thus, in strains bearing a single copy of either *tL(GAG)* or the variant *tL(GAG)C33U* inserted into the *LYS2* gene in the *tL(GAG)Δ* strain, there is no detectable difference in the dose-dependent inhibition by CUC-CCG (Fig. 2C, 2F) or in expression of reporters with two or four inhibitory pairs (Fig. 2E). However, we do find a small, but significant, difference between two strains with a wild type copy of *tL(GAG*) at the native locus, but differing in a second copy of *tL(GAG)* at *LYS2,* one with *C33* (wt) and the other with *U33* (mutant), as we find slightly greater expression of reporters with two or four CUC-CCG pairs in the strain with one copy of *tL(GAG) U33* (Fig. 2D, E). The effects while significant are minor. Thus, we infer that C33 does not substantially impair the function of tRNA^Leu(GAG)^.

### Selection and screening for mutants with a reduction in inhibitory effects of CUC-CCG codon pairs

To examine the mechanisms underlying CUC-CCG inhibition by tRNA^Leu(UAG)^, we constructed strains lacking tRNA^Leu(GAG)^, in which expression of both *URA3* and GFP were reduced by fusion of an N-terminal FN-FN backbone containing CUC-CCG codon pairs to each of the *URA3* and GFP genes (Fig 3A). To further limit URA3 expression, we used a construct in which the *URA3* promoter is missing a UAS that allows upregulation during uracil starvation (Roy et al. 1990). To identify strains suitable for selection, we varied the number of CUC-CCG codon pairs in the FN domains fused to the *URA3* coding sequence and chose strains bearing constructs that significantly limited growth on media lacking uracil; strains with two to four CUC-CCG codon pairs in the FN domains met these criteria. We then performed selections for mutants that grew on media lacking uracil at temperatures from 18°C to 38°C. Mutants that grew on -Ura media were screened for increased expression of GFP/RFP and then subjected to whole genome sequencing. We discuss the analysis of these mutants below.

**Figure 3.**
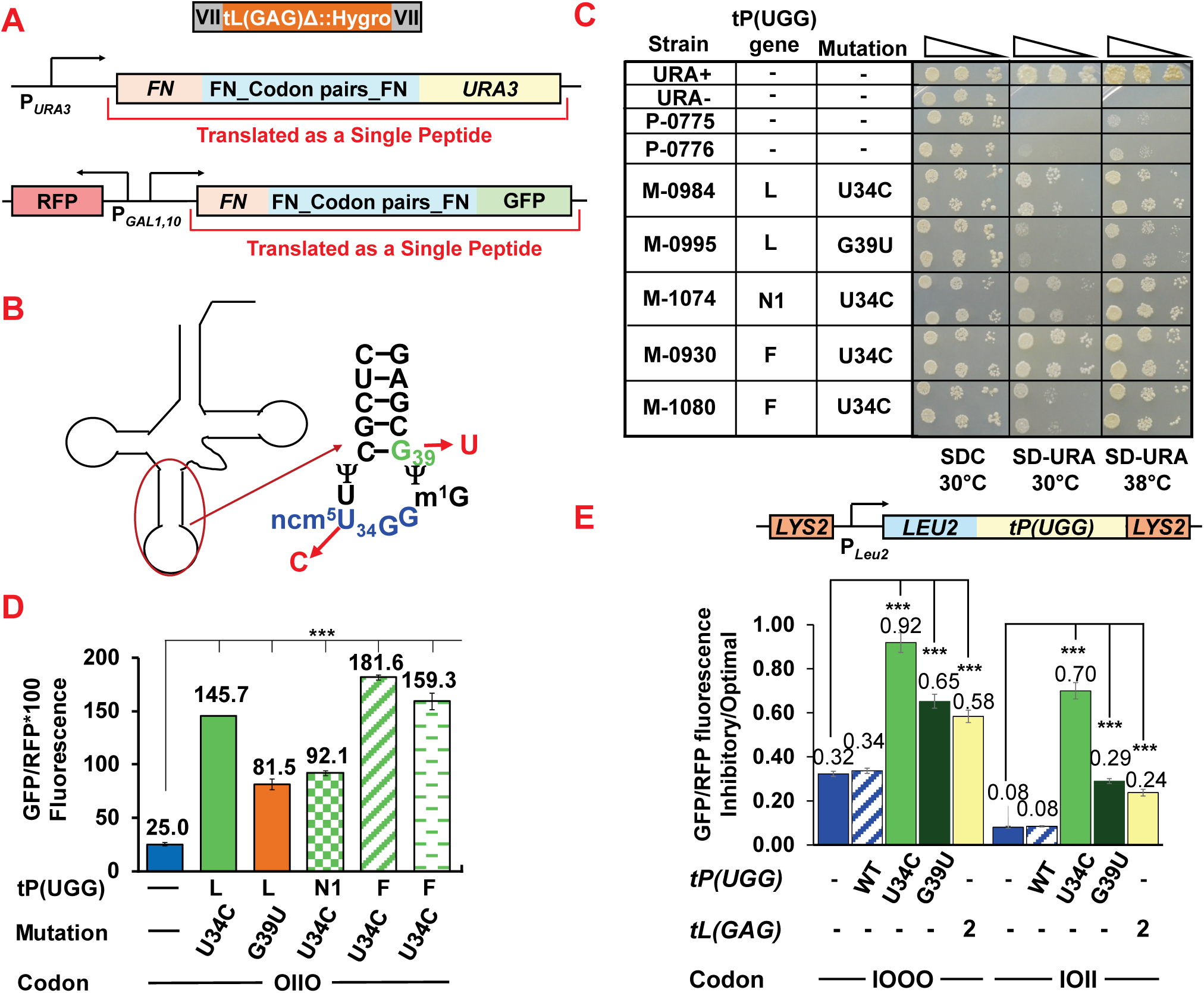
Mutations in tRNA^Pro(UGG)^ that cause increased expression of genes with CUC-CCG affect both the anticodon and closing base of anticodon stem-loop. **A.** Diagram of strain used for selection of mutants that suppress inhibitory effects of CUC-CCG. Mutants were selected on media lacking uracil as expression of *URA3* is limited by separated CUC-CCG codon pairs in same arrangement as used for FN-FN GFP reporter; candidate suppressors were tested for increased expression of GFP/RFP as described previously (Wang et al. 2018; Houston et al. 2022). Candidates that showed a reproducible increase in growth on –Ura media as well as increased GFP/RFP were subjected to whole genome sequencing. **B.** Diagram of tRNA^Pro(CGG)^ showing sites and identities of mutations that resulted in CUC-CCG suppression. The U34C mutation modified the anticodon from UGG to CGG while the G39U mutation affects the closing base of the anticodon stem. **C.** Strains with mutations in the specified *tP(UGG)* genes at the indicated sites exhibit substantially increased growth on media lacking uracil compared to parent strains (P-0775; P-0776), but similar growth on complete minimal media (SDC). Spot tests with serial dilutions of cells from overnight growth. **D.** Strains with mutations in the specified *tP(UGG)* genes at the indicated sites exhibit significantly increased GFP/RFP from a reporter with two inhibitory pairs. *** P value <0.001 compared to parent strain in all cases. **E.** Strains bearing a single inserted copy of mutated *tP(UGG)* (U34C or G39U) suppress inhibition by CUC-CCG more robustly than two copies of tL(GAG). *tP(UGG)* alleles were inserted at *LYS2* as shown. Comparison of expression of GFP/RFP Inhbitory/Optimal ratio from constructs with one (IOOO) or three (IOII) CUC-CCG inhibitory pairs in parent *tL(GAG)Δ* strain (blue), or in strains with a single additional *tP(UGG)* gene inserted at *LYS2* (diagram) (wild type (blue stripe), U34C (lime green) or G39U (forest green)) and in a strain bearing two copies of tL(GAG) (yellow). *** P value <0.001 compared to parent strain.

### Mutations in the anticodon and anticodon stem of tRNA^Pro^ suppress inhibitory effects of CUC-CCG

Six of the suppressor mutants had mutations in four different genes encoding tRNA^Pro(UGG)^ (tP(UGG)); with mutations mapping to either to the anticodon wobble position 34, or to position 39 at the base of the anticodon stem (Fig. 3B). Five of the six mutations alter the anticodon of tRNA^Pro(UGG→CGG)^ converting the tRNA to one that forms W●C base pairs with all three positions of the CCG codon, while one mutation instead alters the closing base of the anticodon stem to produce tRNA^Pro(UGG)^ G39U (Fig.3B; Supplemental Table S1). Strains bearing these mutations exhibit the expected increase in growth on media lacking uracil at 38°C (Fig 3C) and substantial suppression of CUC-CCG inhibition at 30°C, based on expression of a GFP reporter with two CUC-CCG inhibitory codon pairs (Fig. 3D). However, we note that there are substantial differences between the mutants. While all of the mutants grew well (similarly to their parents P-0775 and P-0776) on complete minimal media, they differ in their growth at 30°C or 38°C on media lacking uracil (Fig. 3C), and the mutants differ from each other with respect to GFP/RFP, exhibiting a 2-fold range from 81.5 to 181.6 GFP/RFP from mutants relative to 25.0 from the wild type, corresponding to a 3.3- to a 7.3-fold increase compared to the wild type strain. The strain bearing *tP(UGG)L G39U* appears to be the weakest suppressor, in both assays.

Suppression by the anticodon mutations tRNA^Pro(UGG→CGG)^ was expected as we had previously shown that such a mutant tRNA on a plasmid copy suppressed inhibition by CCG-containing inhibitory codon pairs (Gamble et al. 2016). However, as *S. cerevisiae* encodes 10 copies of *tP(UGG)* (Chan and Lowe 2009; Chan and Lowe 2016), it was unknown if mutation of a single *tP(UGG)* gene would effectively suppress inhibition. In considering the difference in efficacy between the mutants, we speculate that differences in suppression could arise from known differences between these tRNA genes in the sequence of their introns and that such differences may affect their expression levels.

By contrast, there is no clear precedent for the G39U mutation as a wobble suppressor, although it is known that such a mutation almost certainly affects the structure of the anticodon loop or anticodon stem (Grosjean and Westhof 2016). Furthermore, mutations of this type (and elsewhere in the anticodon loop) can promote misreading in vitro (Shepotinovskaya and Uhlenbeck 2013). Thus, it was reasonable to consider that the G39U mutation reduced inhibition by CUC-CCG by improved decoding of the wobble interaction.

To establish that suppression is due to the mutations in *tP(UGG)*, we inserted *tP(UGG)L*, *tP(UGG)L(UGG→CGG)* and *tP(UGG)L(G39U)* genes into the *tL(GAG)Δ* strain by integrating a single copy of the tRNA gene or variant at the *LYS2* locus (Fig 3E). After integrating GFP reporters with zero (OOOO), one (IOOO) or three (IOII) copies of the CUC-CCG inhibitory codon pair into these strains, we compared suppression by a single mutant *tP(UGG)* allele to suppression by two copies of the exact match native *tL(GAG)* to assess the difference in efficacy of the two exact base pairing tRNAs.

We found that a single copy of either *tP(UGG)L(UGG→CGG)* or *tP(UGG)L(G39U)* resulted in increased expression of GFP/RFP from reporters with CUC-CCG inhibitory codon pairs (Fig 3E). The *tP(UGG)L(UGG→CGG)* mutant tRNA is a remarkably efficient suppressor, restoring expression of a reporter with a single CUC-CCG pair from 32% of optimal to 92% of optimal and of a reporter with three inhibitory codons from 8% of optimal to 70% of optimal (Fig. 3E).

By contrast, tRNA^Pro(UGG)^ G39U is a distinct, but notably less efficient suppressor than tRNA^Pro(UGG→CGG)^; when substituted at the same locus, increasing expression of a reporter with a single CUC-CCG pair from 32% to 65%, but only increasing expression of the reporter with three inhibitory codon pairs from 8% of optimal to 27% of optimal (compared to 70% with U34C mutant). This difference in suppression may reflect a difference in effectiveness between the U34C and G39U mutant tRNAs in decoding CCG, although it is possible that the G39U mutation destabilizes the tRNA (Guy et al. 2014) or impairs its general function in translation.

We note that suppression by two copies of *tL(GAG)* is very similar to that by tRNA^Pro(UGG)^ G39U, but much less efficient than that of tRNA^Pro(UGG→CGG)^ (Fig. 2E). Thus, the single copy of tRNA^Pro(UGG→CGG)^ effectively outcompetes the 10 copies of tRNA^Pro(UGG)^ to decode the CCG codon and alleviates most of the inhibitory effects of CUC-CCG, suggesting that A site discrimination against this U●G wobble is very strong in vivo.

To ascertain if the tRNA^Pro^ mutations suppress other inhibitory codon pairs with a Pro CCG codon, we examined expression of inhibitory reporters with CUC-CCG, CUG-CCG, or CGA-CCG, relative to their respective optimal reporters, at amino acid 6 upstream of GFP and at amino acid 100 of GLN4_(1-99)_-GFP reporter (bearing three copies of the codon pair inserted with two amino acid spacers) (Dean and Grayhack 2012; Wolf and Grayhack 2015). The extent of inhibition by these codon pairs varied, with basal expression ranging from 11% to 52% of the optimal reporters (on average 29%) (Supplemental Fig. S6, S7). In all cases, strains with the anticodon mutated tRNA^Pro(UGG→CGG)^ effectively suppressed inhibition, with a median 3-fold increase compared to WT tRNA^Pro^ (Supplemental Fig. S6, S7). By contrast, strains with the anticodon stem mutation tRNA^Pro(UGG)^ G39U displayed weaker, but still distinct, suppression in all but one case, with a median 1.6-fold increase in expression (Supplemental Fig. S6, S7). Although the tRNA^Pro(UGG)^ G39U did not suppress the CUC-CCG inhibitory pair at amino acid 100, although it did strongly suppress CUC-CCG at amino acid 6 (3.8-fold). Thus, the effects of tRNA^Pro^ ^(UGG)^ G39U are likely context dependent, although in five of the six cases, this mutant tRNA did suppress inhibition by other codon pairs containing Pro CCG.

### Mutations in several large subunit proteins and a chaperone for a large subunit protein suppress inhibition by CUC-CCG

Among the suppressor mutants, we identified six strains with mutations in one of two redundant copies of large subunit ribosomal proteins, and one strain with a mutation in an essential chaperone for a large subunit ribosomal protein (Supplemental Table S1). All of the ribosomal protein mutations are likely to produce a nonfunctional copy of the protein as they bear nonsense or frameshift mutations (Supplemental Table S1). All four re-tested suppressor strains bearing *RPL* mutations grow substantially better than their parents on media lacking uracil at 30°C (near their selection temperature) (Fig 4A). Two strains, with *rpl4A* and *rpl7A* mutations, also grow much better than their parent on SD-Ura at 37°C, and all four suppressors unexpectedly grow very poorly on YPD, but not SDC, medium (Fig 4A). In addition, six *rpl* suppressors strains exhibit similar ∼2-fold increase in expression of an FN-FN GFP reporter with two CUC-CCG inhibitory pairs. Similarly, the strain with a mutation in chaperone *BCP1* also grows better than its parent on media lacking uracil and show increased GFP (Supplemental Fig. S8).

**Figure 4.**
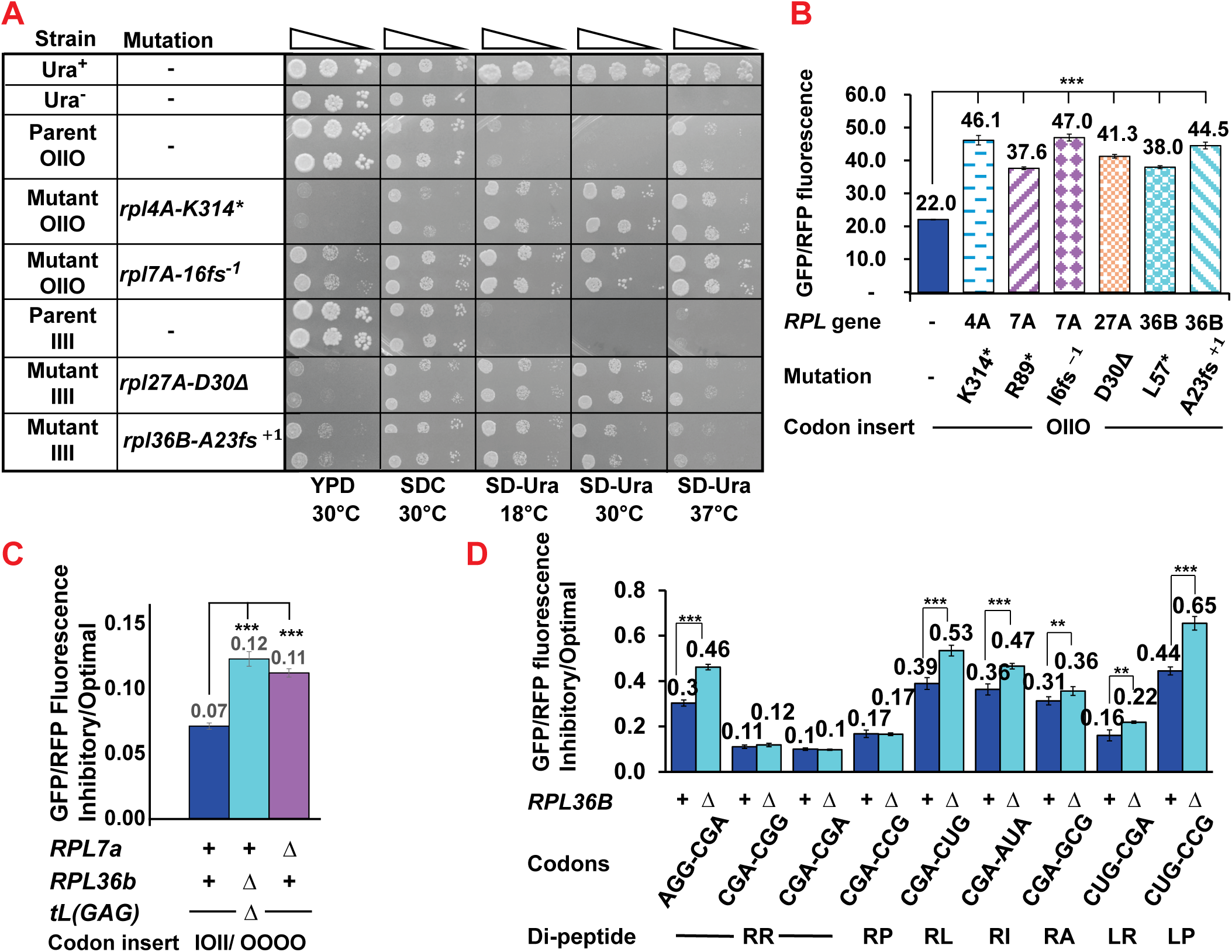
Mutations that suppress CUC-CCG inhibition map in large subunit ribosomal protein genes. **A.** Suppressor strains with mutations in each of four genes encoding large subunit ribosomal proteins exhibit increased growth on media lacking uracil and decreased growth on rich (YPD) media. Spot test showing growth of serially diluted yeast from overnight growth in liquid media. **B.** Strains with the indicated mutations in ribosomal protein genes exhibit increased GFP/RFP relative to the parent strain from an integrated GFP reporter with two CUC-CCG inhibitory pairs. **C.** Reconstructed strains with deletions of *RPL7A* or *RPL36B* exhibit a significant increase in expression of reporters with three CUC-CCG inhibitory pairs compared to expression of reporters with optimal pairs. GFP reporters were integrated into indicated strains bearing a covering *RPL7A URA3 Cen* or *RPL36B URA3 Cen* plasmid, followed by growth on FOA to select for loss of the plasmid, and then analysis of GFP/RFP by flow. Wild type (blue), *rpl36BΔ* (cyan), *rpl7AΔ* (purple). **D.** Six of nine tested inhibitory pairs are suppressed in strains bearing an *RPL36B* deletion. The ratio of GFP/RFP from reporters with three inhibitory pairs inserted in an IOII arrangement to GFP/RFP from reporters with synonymous optimal pairs is shown in parent *tL(GAG)Δ* (blue) and the *tL(GAG)Δ rpl36BΔ* mutant (cyan). In each case, the reporters were transformed into strains bearing chromosomal *rpl36BΔ*::kan and the covering plasmid *RPL36B URA3 CEN*, followed by selection on FOA and purification, prior to analysis of GFP/RFP by flow. *** P value <0.001.

To determine if reduction in the ribosomal protein is responsible for suppression, we constructed strains in which each of two *RPL* genes (*RPL7A* and *RPL36B*) were deleted, but the health of the strains was maintained by a plasmid borne copy of the relevant *RPL* gene. To examine effects of deleting *RPL7* or *RPL36B* on inhibition by CUC-CCG, we inserted reporters with all optimal codon pairs (OOOO) or three inhibitory codon pairs (IOII), then selected for loss of the plasmid copy of the *RPL* gene by growth on FOA and measured expression of GFP/RFP (see (Wolf and Grayhack 2015)). Strains bearing either the *rpl7AΔ* or the *rpl36BΔ* mutation strongly suppress inhibitory effects of CUC-CCG as the ratio of expression of inhibitory/optimal reporters in the mutants is much greater than in the wild type strain, from 0.07 (wt) to 0.12 *(rpl36BΔ*), a 1.7-fold increase in *rpl36BΔ* and from 0.07 to 0.11 (*rpl7AΔ*), a 1.5-fold increase in *rpl7AΔ*. (Fig. 4C, Supplemental Fig. S9).

It seems likely that suppression by the mutations in the *RPL* (and *BCP1*) genes is mediated by a reduction in ribosome concentration, for two reasons. First, Woolford and colleagues established that deletion of *RPL7A* results in reduced levels of 60S subunits (Jakovljevic et al. 2012). Second, the identity of other suppressors provides support for the idea that a reduction in ribosomes or 60S subunits is likely responsible for suppression. Thus, we also isolated four suppressors with mutations in genes that mediate ribosome or large subunit assembly factors (two in *NOG2*, one each in *NOP4* and *RSA1*) (Massenet et al. 2017; Klinge and Woolford 2019) and two mutations in *RPA190* (the largest subunit of RNA polymerase I) (Supplemental Table S1).

We considered the possibility that CUC-CCG inhibition is mediated by a combined effect of Hel2 or Mbf1 with another regulator as each of these proteins recognize collided ribosomes, which are found at CUC-CCG codon pairs in yeast (Meydan and Guydosh 2020). Hel2 (homolog of human ZNF598) is one of the most direct sensors of collided ribosomes, triggering NGD decay (Juszkiewicz et al. 2018; Ikeuchi et al. 2019), but there may be additional mechanisms to trigger NGD. For instance, Veltri et al. (Veltri et al. 2022) found that deletion of both *SYH1* and *HEL2* resulted in synergistic enhancement of expression of a reporter with CGA codon repeats. They concluded that Hel2 drives one branch of NGD, while Syh1 and Smy2 drive an independent NGD pathway (Veltri et al. 2022). Mbf1 (homolog of human EDF1) interacts directly with collided ribosomes, prevents frameshifting at CGA-CGA codon pairs, and mediates induction of the ISR, most likely through effects on GCN1 (Wang et al. 2018; Juszkiewicz et al. 2020a; Sinha et al. 2020; Kim et al. 2024). As its human homolog Edf1 is responsible for recruitment of translational repressors GIGYF2 and eIF4E2 (Juszkiewicz et al. 2020a; Sinha et al. 2020), we considered that Mbf1 might have unknown functions in yeast and coordinated effects with other proteins. To test these ideas, we selected a set of suppressor mutations in a *hel2Δ tL(GAG)Δ* background and another set in an *mbf1Δ tL(GAG)Δ* background and subjected these mutants to whole genome sequencing to identify candidate mutations (Supplemental Tables S2, S3).

In the *hel2Δ tL(GAG)Δ* strain, we identified three mutations in large ribosomal subunit proteins (*RPL36B L343\**; *RPL31A–*intron -2 from 3’ splice site, *RPL10 H51L*) and a mutation in *RPA190*, while in the *mbf1Δ tL(GAG)Δ* strain we identified a frameshift mutation in *CGR1* (involved in rRNA processing for the 60S subunit). We also identified nonsense mutations in *PXR1* (involved in rRNA and snoRNA maturation) in both *mbf1Δ* and *hel2Δ* backgrounds, as well as mutations in *SCH9* (See below) (Supplemental Tables S2, S3). Thus, while we did not find evidence of a second pathway working with either Hel2 or Mbf1, (although this is not conclusive proof that such a pathway does not exist), we did reinforce the prevailing evidence that the concentration of either large ribosomal subunits or ribosome is crucial to inhibition, even in the absence of two molecules that bind to collided ribosomes.

### Loss of large subunit protein Rpl36B suppresses inhibition by several but not all inhibitory codon pairs

To find out if loss of Rpl36B affects other inhibitory codon pairs, we examined expression of reporters with each of nine different inhibitory codon pairs, relative to reporters with their corresponding synonymous optimal pairs. We find that the *rpl36BΔ* mutant suppresses the inhibition by all three inhibitory pairs with a Leu CUG codon in either the first or second position (CGA-CUG, RL; CUG-CGA, LR; CUG-CCG, LP) and by three other inhibitory pairs (AGG-CGA, RR; CGA-AUA, RI; CGA-GCG, RA) (Fig. 4D, Supplemental Fig. S10). We find that *rpl36BΔ* results in a 1.47-fold increase in expression of reporter with a CUG-CCG inhibitory pair (0.65 in *rpl36BΔ* compared to 0.44 in WT); for the six codon pairs suppressed by *rpl36BΔ*, the fold suppression ranges from 1.14 for the CGA-GCG (RA) to 1.52 for AGG-CGA (RR), with an average of 1.4.

Inhibition by three pairs (CGA-CGA, RR; CGA-CGG and CGA-CCG, RP) is not effectively suppressed by the *rpl36B*Δ (Fig. 4D). This is a particularly surprising observation as these are the three pairs for which it is known that ribosome collisions mediate their effects, acting via Hel2, the No Go decay pathway and the ribosome quality control complex (Simms et al. 2017a; Wang et al. 2018). Thus, it is surprising that loss of this large subunit protein does not suppress their inhibitory effects.

### CUC-CCG inhibition depends upon ribosome abundance on the specific transcript

To determine the specific requirements for suppression of CUC-CCG inhibition, we assessed the effects of loss of other ribosomal proteins as well as the effects of a reduction in ribosomes on a single transcript. To determine if suppression of CUC-CCG inhibition is due to mutation of the particular large subunit proteins identified in our selection or to reduced abundance of the large subunit or ribosomes in general, we examined the effect of deleting *RPL1B* on CUC-CCG inhibition. Mutations in *RPL1B* and its deletion had previously been shown to suppress CGA-CGA inhibition (Letzring et al. 2013), and to reduce both ribosome abundance and ribosome collisions (Simms et al. 2017b). We found the ratio of inhibitory to optimal GFP/RFP increased significantly in the *rpl1BΔ* mutant for both tested constructs with one and three CUC-CCG inhibitory pairs (IOOO and IOII) (Fig. 5A); 1.3-fold for the reporter with a single inhibitory pair (0.38 compared to 0.29) and 1.7-fold for the reporter with three inhibitory pairs (0.11 compared to 0.07). As we had previously observed with the *rpl36BΔ* mutant, GFP/RFP increased for all reporters, including the optimal reporter, in the *rpl1BΔ* mutant, but the increase in GFP/RFP from reporters with inhibitory pairs was proportionally greater (Supplemental Fig. S11). Thus, we infer that deletion of *RPL1B* suppresses CUC-CCG inhibition, an argument that suppression is due to loss of either large subunits or ribosomes, but not the particular ribosomal protein.

**Figure 5.**
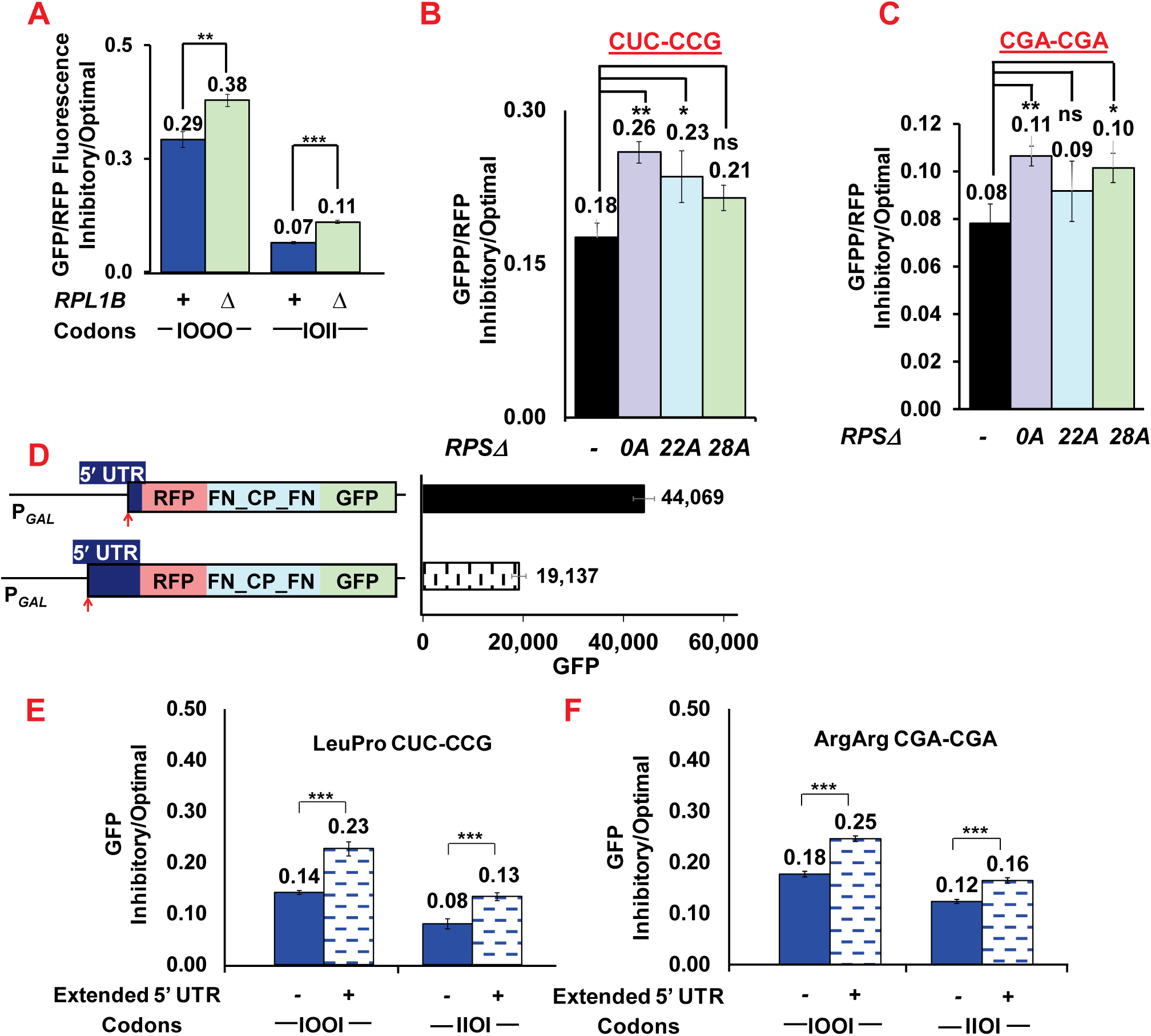
Effects of CUC-CCG depend upon ribosome concentration or density on an individual transcript. **A.** Deletion of *RPL1B* encoding a large subunit protein suppresses inhibition by CUC-CCG. Ratio of GFP/RFP expression of reporters with indicated CUC-CCG inhibitory pairs to that of reporters with optimal pairs in wild type and *rpl1BΔ* strains. In each case, reporters were integrated into strains bearing a *URA3 CEN RPL1B* plasmid, then streaked on FOA to promote loss of the plasmid, followed by analysis of GFP/RFP. **B., C.** Deletion of any one of three small subunit proteins suppresses inhibition by CUC-CCG (B) and by CGA-CGA (C). Ratio of expression of reporters with inhibitory pairs to that of reporters with optimal pairs in wild type and indicated *rpsΔ* mutants B.-CUC-CCG reporters; C.CGA-CGA reporters. **D.** A 360 nucleotide 5’ UTR from *YDL224C* results in reduced GFP compared to GFP from an otherwise identical reporter with a 10 nucleotide 5’ UTR from *PGK1*. **E., F.** Inhibition by either CUC-CCG (E) or CGA-CGA (F) is suppressed in the reporter with the 360 nucleotide 5’UTR from *YDL224C* compared to otherwise identical reporters in with a 10 nucleotide 5’UTR from *PGK1*. Ratio of expression of reporters with inhibitory pairs to that of reporters with optimal pairs is shown.

To determine if loss of small subunit ribosomal proteins can suppress CUC-CCG inhibition, we assayed expression of reporters with all optimal codons and with three inhibitory codons in strains bearing a single deletion of redundant small subunit proteins. In three different mutants, we observed an increase in inhibitory to optimal GFP/RFP (IOII/OOOO) from 0.18 (wild type) to 0.21, 0.23, or 0.26 in the *rps* mutants (1.2 to 1.4 fold) (Fig. 5B), although only the largest of these increases was statistically significant. As inhibition by CGA-CGA codon pairs is mediated by ribosome collisions and thus expected to be affected by ribosome abundance, we also examined the effects of the same three deletions on CGA-CGA inhibition. Again, we found a relative increase in expression of CGA-CGA containing reporters in all three mutants, although the results were only statistically significant in the same single *rps0AΔ* strain (Fig. 5C). We infer that an overall reduction in ribosome concentration or loss of small subunits can suppress the effects of either CUC-CCG or CGA-CGA, but loss of small subunit proteins is unlikely to be as effective at suppression as loss of the large subunit proteins.

To determine if the local or global reduction in ribosomes was required for suppression, we examined effects of reducing ribosome load on a single transcript using a long 5′ UTR, as described by Simms *et al*. (Simms et al. 2019). We compared expression of GFP from two RFP-FN-GFP reporters differing only in their 5′UTRs, with one 5’UTR bearing the 10 bases upstream of *PGK1* and a second 5’ UTR bearing the 360 nucleotide 5’UTR from *YDL224C*. As expected, we found that expression of GFP from the reporter with the 5′UTR from *YDL224C* was reduced to 43% compared to that from the reporter with the 5’ UTR from PGK1 (19,137 compared to 44,069) (Fig. 5D).

We assessed the magnitude of CUC-CCG inhibition in these two reporters, by comparing expression in the *tL(GAG)Δ* strain of reporters with two and three inhibitory CUC-CCG codon pairs to expression of a synonymous optimal reporter (Fig. 5E, Supplemental Fig. S.12). In each case, the inhibitory effect of CUC-CCG was significantly greater in the transcript with the short 5’ UTR (compare 0.14 for two inhibitory pairs with the short 5’ UTR compared to 0.23 for two inhibitory pairs with the long 5’ UTR), a 1.6-1.7-fold effect. Moreover, as expected, we observe the same phenomenon with CGA-CGA in both strains (Fig. 5F; Supplemental Fig.S13). Thus, the inhibitory effects of CUC-CCG, like those of CGA-CGA, depend upon the ribosome density on the particular mRNA, consistent with the idea that inhibition depends upon ribosome abundance and collisions.

### The TORC1 regulator Sch9 modulates inhibition by CUC-CCG

Six of the suppressor mutants carried mutations in *SCH9*, the TORC1 regulated AGC-family vacuolar kinase which is itself one of the two major effectors by which TORC1 regulates cell growth (Huber et al. 2009) (Fig. 6A, Supplemental Table S1). Sch9 protein is an 824 amino acid protein bearing domains at its N terminus and adjacent to the N terminus (C2) that are responsible for its localization to the vacuole, while its C terminal region mediates the critical kinase function and is itself targeted by numerous kinases (Fig. 6A) (Caligaris et al. 2023). Sch9 is best known as a critical regulator of the translation machinery, regulating synthesis of rRNA by controlling RNA polymerase I recruitment to rDNA promoters, regulating expression of ribosomal proteins and ribosome biogenesis factors by phosphorylation and inactivation of transcriptional repressors, and regulating tRNA expression by phosphorylation of the master regulator Maf1 (Gutierrez-Santiago and Navarro 2023). The suppressors bear *SCH9* mutations mapping in the kinase domain from R509 to K682 (Fig. 6A; Supplemental Table S1).

**Figure 6.**
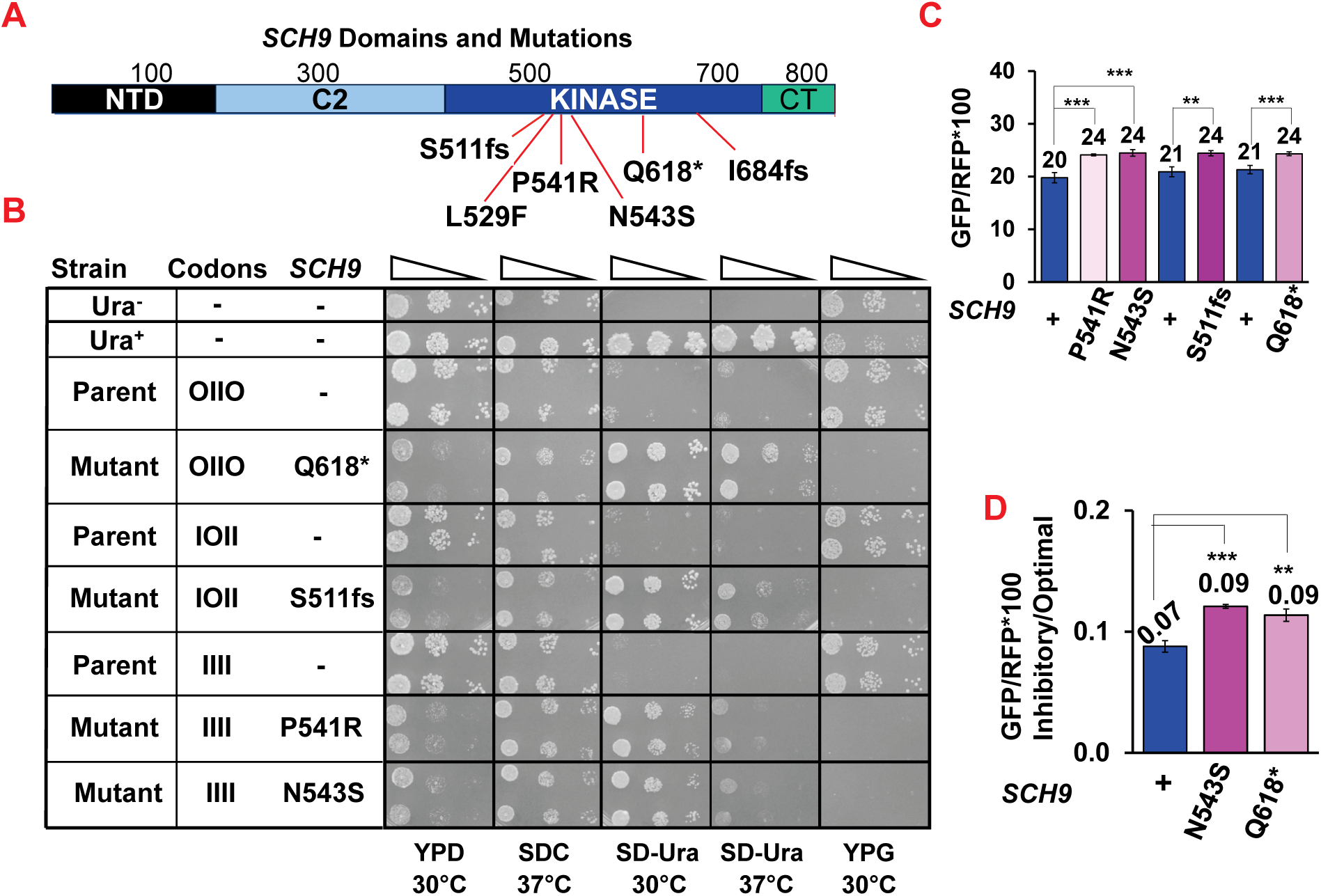
A role for Sch9. a TOR regulated kinase, in CUC-CCG inhibition. **A.** Diagram of Sch9 protein domains, showing locations and identity of mutations identified as suppresors of CUC-CCG inhibition. **B.** Strains with indicated mutations in *SCH9* exhibit substantially increased growth on media lacking uracil compared to parent strains. As expected strains with *sch9* mutations fail to grow on media with glycerol as the carbon source and grow poorly on rich media (Peterson and Liu 2021). **C.** Strains with indicated mutations in *SCH9* exhibit significant but small increases in GFP/RFP expression of a reporter with two inhibitory pairs. **D.** Reconstructed strains bearing wild type, *sch9 N543S* and *sch9 Q618\** suppress CUC-CCG inhibition. The ratio of GFP/RFP from reporters with 3 CUC-CCG inhibitory codon pairs compared to that from reporters with all optimal pairs is reported. *** P value <0.001

Strains bearing these *sch9* mutations exhibit growth properties consistent with suppression of CUC-CCG inhibition beyond the selection temperature (18°C) and indicative of reduced function of Sch9. Four tested strains bearing mutations in *SCH9* (Q618*, S511fs, P541R and N543S) all grow much better than their respective parents on media lacking uracil at 30°C, with two strains (Q618* and S511fs) also growing well at 37°C (Fig. 6B). The mutants also display two phenotypes expected for mutants in which Sch9 function is compromised: they all grow considerably more slowly than their parents on rich media (YPD) (Fig. 6B) as expected for a defect in sensing nutrient abundance, and also fail to grow on glycerol media, a known phenotype of *sch9* null alleles (Fig. 6B) (Peterson and Liu 2021). All four tested mutants exhibited modest increased expression of GFP reporter with two CUC-CCG inhibitory codon pairs at 20°C (Fig. 6C), as expected.

To determine if the *sch9* mutations were causative for reduced inhibition by CUC-CCG, we reconstructed two mutations (N543S and Q618*) in the *tL(GAG)Δ* strain, and then inserted FN-GFP reporters bearing either four optimal (OOOO) or three inhibitory CUC-CCG codon pairs (IOII). Examination of GFP/RFP expression at 30°C showed only a minimal increase in expression from the optimal constructs (1.1 fold), but a significantly larger increase in both reconstructed mutants from GFP reporters with three inhibitory pairs (Supplementary Fig. S14). The ratio of inhibitory to optimal expression rose from 0.07 in the wild type to 0.09 in each of the mutants, a 1.3-fold increase in expression, with high statistical significance (Fig. 6D). Thus, we infer that the *sch9* mutations are responsible for reduced inhibition, albeit a relatively small effect.

### Sch9 also modulates inhibition by CUC-CCG in the absence of Hel2 or Mbf1

In both the *hel2Δ* and *mbf1Δ* strains, we identified suppressor strains with mutations in *sch9* in the suppressor strains, three of the eleven suppressors in the *hel2Δ* background and one of the four suppressors in *mbf1Δ* background bear mutations in *SCH9* and no other candidate mutations. Strains bearing these mutations grow significantly better than their parents on media lacking uracil, as expected if the mutants suppress CUC-CCG inhibition of *URA3* expression (Supplemental Fig. S15). They also grow poorly on rich media and do not grow on glycerol as expected if Sch9 function is compromised. All of the strains with *sch9* mutations also exhibit increased expression of the GFP reporter with 2 inhibitory codon pairs (Supplemental Fig. S16). Thus, we infer that Sch9 does mediate inhibition by CUC-CCG, likely in a manner that is not dependent upon Hel2 or Mbf1 and thus not dependent upon the standard NGD pathway.

## DISCUSSION

One major finding that emerges from these studies is that inhibitory codon pairs likely regulate gene expression in response to the metabolic status of the cell, which is communicated by changes in the concentration of ribosomes. For the Leu-Pro CUC-CCG codon pair, we conclude first that the ribosome concentration is the critical factor in its inhibitory effects, based on findings that CUC-CCG inhibition is relieved by numerous mutations expected to reduce ribosome concentration, by explicit depletion of any of three large ribosomal subunit proteins or one small ribosomal subunit protein, and by a reduction in ribosome density on a single transcript. We deduce that inhibition by most inhibitory codon pairs is dependent upon ribosome concentration, as this is known for the NGD-activating pairs CGA-CGA, CGA-CGG and CGA-CCG (Simms et al. 2017b; Simms et al. 2019). We further infer that other inhibitory pairs are regulated by the concentration of ribosomes, as explicit depletion of one large ribosomal subunit protein (a suppressor of CUC-CCG) also suppresses inhibition by six of six other non-NGD inhibitory pairs. Thus, there is evidence that 10 of the 17 pairs are responsive to ribosome concentration. We judge that the magnitude of inhibition is responsive to the metabolic state of the cell as we find six mutations in *SCH9* that suppress inhibitory effects of CUC-CCG. Sch9, an AGC family kinase that is a direct target of TORC1 and one of two major mediators of TORC1 regulation (Urban et al. 2007), is known to regulate production of tRNAs, ribosomal proteins and ribosomal RNAs (Gutierrez-Santiago and Navarro 2023) and is inactivated in starvation conditions (Urban et al. 2007). Thus, suppression of CUC-CCG inhibition by impaired Sch9 function connects the regulation of gene expression by inhibitory codon pairs, with the ribosome concentration and the metabolic state of the cell.

A second inference that emerges from this study is that tRNA^Leu(GAG)^ and tRNA^Leu(UAG)^ differ substantially from each other in their function in the P site of the ribosome, and thus, in wild type yeast the inhibitory effects of any occurrence of the CUC-CCG pair can have different consequences, dependent upon the actual tRNA accepted to decode the CUC. This inference is based on the finding that tRNA^Leu(GAG)^ and tRNA^Leu(UAG)^ compete to decode the CUC codon but have different effects on the inhibition conferred by CUC-CCG. As we have argued previously that codon pair inhibition most likely involves a 5’ codon in the P site that influences the rate of acceptance of tRNA for the 3’ codon in the A site (Gamble et al. 2016), (based on tRNA suppression data and accumulation in the P-A site of the ribosome, and as shown directly for CGA-CGA (Tesina et al. 2020)), we infer that the tRNA^Leu(GAG)^ and tRNA^Leu(UAG)^ have functional differences in the P site of the ribosome. We estimate that a single CUC-CCG codon pair fully decoded by tRNA^Leu(UAG)^ reduces expression to 0.33 (Fig. 2) while a single CUC-CCG codon pair fully decoded by tRNA^Leu(GAG)^ would reduce expression to ∼0.7 (Supplemental Fig. S4), and based on measured inhibition in wild type yeast, we infer that ∼50% of the CUC-CCG pairs are decoded by the U●C wobble interacting tRNA^Leu(UAG)^. As there is clear evidence that many codons can be decoded by different tRNA species (Johansson et al. 2008) and that P site interactions are sensitive to wobble decoding tRNAs (Letzring et al. 2010; Tunney et al. 2018; Tesina et al. 2020), our results provide a clear biological example of the impact that such competition can have.

We deduce that there are major differences in A site discrimination of different wobble interacting tRNAs. Surprisingly we find that a single copy of the *tP(UGG→CGG)* gene in a cell with 10 other genes bearing the wild type wobble sequence *tP(UGG)*, nearly completely suppresses inhibitory effects of even three copies of the CUC-CCG codon pair (from 0.08 to 0.70, Fig. 3), an argument that the ribosome overwhelmingly prefers the exact base pairing tRNA and must reject the wobble decoding tRNA. The single copy *tP(UGG→CGG)* mutant tRNA is a strong suppressor of all three tested inhibitory codon pairs with a Pro CCG codon (CUC-CCG, CUG-CCG, and CGA-CCG), and suppresses inhibition by these pairs at amino acid 6 or 100, sites which differ in the mechanism of inhibition. By contrast, we find that even two copies of the exact match *tL(GAG)* in a cell compete relatively poorly with the three copies of *tL(UAG)*. Thus, it appears that the ribosome does not strongly discriminate against the U●C mismatch required for tRNA^Leu(UAG)^ to decode CUC. In this context, it may be relevant that tRNA^Leu(UAG)^ is the only tRNA in *S. cerevisiae* with an unmodified U34 residue (Randerath et al. 1979; Johansson et al. 2008), as there is abundant evidence that tRNA modifications modulate interactions between codon and anticodon in the ribosome, particularly wobble decoding (see (Kothe and Rodnina 2007)).

While the tRNA^Pro(UGG)^ G39U suppressor could affect the interactions between the anticodon of the tRNA and the Pro CCG codon, it might exert its effects by other mechanisms. On the one hand, as part of the proximal extended anticodon, the 31-39 base pair stabilizes the helical conformation of the anticodon stem and likely affects the structure of the anticodon loop, which could directly affect codon-anticodon interactions (Grosjean and Westhof 2016). On the other hand, the essential kinetic discrimination between cognate and near-cognate tRNAs (Gromadski and Rodnina 2004) can be modulated in different ways as several mutations distant from the anticodon stem-loop affect tRNA fidelity (Cochella et al. 2007; Schmeing et al. 2011; Schrock et al. 2025). For instance, the Hirsch suppressor G24A in the D-arm of tRNA^Trp^ stimulates GTP hydrolysis by EF-Tu (Cochella and Green 2005), likely due to formation of a hydrogen bond that stabilizes the distorted tRNA conformation (Schmeing et al. 2011), while an A9C mutation in tRNA^Trp^ also stabilizes a distorted tRNA conformation likely due to increased flexibility of the tRNA (Schmeing et al. 2011). Thus, it would be of interest to examine the mechanism by which G39U suppresses inhibitory effects of CUC-CCG and to determine if its effects are also seen in the absence of the CUC codon.

While our evidence points strongly to the idea that inhibition mediated by CUC-CCG, similar to that of CGA-CGA, is likely mediated by ribosome collisions, we note that there are clearly major differences in the mechanisms that mediate effects of different inhibitory codon pairs. We know that inhibition by the three NGD-activating pairs, CGA-CGA, CGA-CGG, CGA-CCG, is suppressed by mutations affecting Asc1 (RACK1) or Hel2 (ZNF598), and that +1 frameshifting at these pairs is revealed by mutations affecting Mbf1; by contrast, none of nine other tested inhibitory codon pairs (including CUC-CCG) displayed these phenotypes. Here we found that CUC-CCG inhibition, as well as that of six other tested pairs, was suppressed by deletion of one of two copies of genes encoding Rpl36 while, surprisingly, none of the NGD-activating codon pairs was suppressed by this mutation. Thus, there are likely differences between translation of different inhibitory pairs that trigger different quality control mechanisms. We still do not know the factors that cause degradation of mRNA or prevent continued translation by collided ribosomes at the CUC-CCG codon pair or at any of the other non-NGD codon pairs. While we did not obtain mutations that point to a specific system, we know that there are several translation rate-dependent mRNA degradation systems (Muller et al. 2025). One possibility is that inhibitory effects of CUC-CCG are mediated through multiple redundant systems, encoded by multiple redundant genes (like Cue2 or rRNA). An alternative is that the system that responds to CUC-CCG codon pairs cannot be easily mutated as it might be essential for life, such as at the ribosome itself.

## MATERIALS AND METHODS

### Strains, Plasmids, oligonucleotides and Gene Blocks

Strains, plasmids, oligonucleotides and gene blocks used in this study are listed in Supplemental Tables S4-S6; yeast strains with suppressor mutations are listed in Supplemental Tables S1, S2 (*hel2Δ* parent) and S3 (*mbf1Δ* parent). BY4741 was the parent for yeast strains used in these studies (Open Biosystems). Strains bearing deletions of *tL(GAG)* or other yeast genes were constructed by standard methods using the genomic deletion collection (Giaever et al. 2002) or plasmid cassettes bearing resistance markers (Wach et al. 1994; Goldstein and McCusker 1999; Gueldener et al. 2002). The plasmid EEVN391, a derivative of the RNA-ID reporter (Dean and Grayhack 2012), constructed by insertion of gene block EP14 into EagI cut ELB798 vector, resulted in an FN-FN-GFP construct allowing insertion of codon pairs into FN loops with 32-59 codons between each insert; EEVN391 sequence from Plasmidsaurus is reported in Supplemental Figure S17. Integrating vectors bearing wild type or variants of tL(GAG) or tP(UGG) were constructed using Gibson cloning into BamH1 cut EAH118-2, an integrating vector with homology to *LYS2* on both ends and *LEU2* gene for selection; the sequence of EAH118-2 is reported in Supplemental Figure S17. Derivatives of EEVN535 used in selections for suppressors contain different numbers of CUC-CCG inhibitory codon pairs in the FN domains upstream of URA. EEVN535 contains sequences homologous to the *URA3* promoter (to -508), with a 16 base pair deletion of the *URA3* UAS from -143 to -158 (Roy et al. 1990), a fusion at amino acid 5 of *URA3* coding sequence to sequences with homology to FN domains from EEV391, the remaining *URA3* coding sequence and a copy of *S. pombe HIS5* gene downstream of *URA3*. Fusion of FN domains with zero to four inhibitory codon pairs modulates expression of *URA3*, resulting in Ura+ to Ura-phenotypes. The sequence of EEVN535 is reported in Supplemental Figure S17.

The plasmid EBB024 is a modified RNA-ID reporter, expressing a FLAG-RFP-GFP fusion protein, set up to facilitate insertion of sequences between RFP and GFP, either by standard restriction sites or by Gibson cloning. The sequence is reported in Supplemental Figure S17, EBB024 was modified with gbEP021 to include a 371 base 5’UTR from YDL224C directly upstream of RFP at the NheI site.

### Selection of mutants that suppress CUC-CCG inhibition

Selection for mutants that suppress CUC-CCG inhibition was similar to previous selections for suppressor that allow frameshifting at CGA-CGA codon pairs (Wang et al. 2018; Houston et al. 2022), in that 2 million cells were plated on SD-Uracil plates, which were incubated at ranges of temperatures from 18-38°C for up to 20 days, counted apparent mutants over 20 days. Putative mutants were streaked on SD-Ura and grown at the selection temperature, then grown in liquid YPD media and saved at -80°C. The mutant was streaked from the save, grown on YPD plates, then three colonies were patched on YPD, followed by overnight growth in YP with raffinose and galactose and analysis by flow cytometry. Mutants with a 1.2-fold increase in GFP/RFP were considered likely to contain suppressor mutations, were subjected to second round of flow analysis and growth analysis on -Ura media by spot testing (see below).

### Identification of mutations in suppressor strains by whole genome sequencing

Mutations were identified by whole genome sequencing and mapped to the yeast genome by methods described previously (De Zoysa et al. 2024). Whole genome sequencing was performed by Cornell Genomics Facility at read depths of 20-80 reads per nucleotide.

### Construction of strains bearing deletions of *RPL7A* and *RPL36B*

Deletions of *rpl7A* and *rpl36B* were generated in the diploid BY4743. The *tL(GAG)Δ* parent strain (YBB0242) was transformed with either *RPL7A URA3 CEN* plasmid or *RPL36B URA3 CEN* plasmid, followed by integrative transformation of the *rpl7AΔ::KanR* or *rpl36::KanR* into the strain bearing its complementary plasmid. Chromosomal deletions of the relevant *rpl* gene were confirmed by PCR. Similarly, deletion of *rpl1B* was performed in a strain bearing a *CEN URA3 RPL1B* plasmid.

### Analysis of Yeast Growth

Yeast strains were grown overnight at 30°C, diluted in sterile water to obtain and OD600 of 0.5, followed by serial dilutions in sterile water, with 2 μl spotted on the indicated plates and incubate at the indicated temperatures.

### Flow Cytometry and RT-qPCR

Analysis of GFP and RFP by Flow cytometry and assessment of p-values for this data were performed as described in Houston et al. (Houston et al. 2022). Analysis of mRNA levels by RT-qPCR and assessments of significance were performed as described previously (Gamble et al. 2016; Wang et al. 2018). P values were calculated using a one-tailed or two-tailed homoscedastic t test in Excel, as indicated in the source data for relevant figures.

For strains bearing deletions of RPL genes (7A, 36B, 1B), reporters were integrated into strains bearing the relevant *URA3 CEN* complementing *RPL* plasmid; transformants were purified, then streaked onto FOA to promote loss of the plasmid. Single colonies picked from FOA were grown in galactose-containing media, followed by analysis of GFP and RFP.

## SUPPLEMENTAL MATERIAL

Supplemental material is available for this article.

## ACKNOWLEDGEMENTS

We thank Eric Phizicky for discussions of the science and comments on the manuscript, Monika Tasak for strains beaing *tL(UAG)* deletions, Alayna Hauke and Eric Phizicky for *LYS2* vector and Justin Fay for guidance on mapping mutations. We thank Cornell University Genomics Center for high-throughput sequencing. This work was supported by National Institutes of Health (NIH) grant R01 GM118386 to E.J.G. L.B.H. was also supported by an NIH T32 Training Grant in Cellular, Biochemical and Molecular Sciences (GM68411).

## Abbreviations

NGD: No-Go decay
ISR: Integrated Stress Response pathway

---

**Supplemental Figure S1. Effects of *gcn1Δ hel2*Δ on inhibition by CUC-CCG.** GFP/RFP fluorescence and GFP/RFP mRNA from reporters with Leu-Pro UUG-CCA (Optimal) at all 4 positions and with CUC-CCG (inhibitory) pairs at three positions in wild type and *gcn1Δ hel2*Δ strains. (Data used in Fig. 1C).

**Supplemental Figure S2. Effects of *gcn1Δ hel2*Δ on inhibition by CGA-CGA.** GFP/RFP fluorescence and GFP/RFP mRNA from reporters with Arg-Arg CGU-CGU (Optimal) at all three positions and with CGA-CGA (inhibitory) pairs at three positions in wild type and *gcn1Δ hel2Δ* strains. The second codon pair in both constructs is an optimal Leu-Pro pair UUG-CCA. (Data used in Fig. 1D).

**Supplemental Figure S3. Inhibitory effects of CUC-CCG are observed in strains lacking tRNA^Leu(GAG)^.** Comparison of GFP/RFP fluorescence (protein) and mRNA from reporters with indicated Leu-Pro optimal (UUG-CCA) and inhibitory (CUC-CCG) codon pairs in the *tL(GAG)Δ* strain.

**Supplemental Figure S4. Reduction in copy number of *tL(UAG)* genes results in increased expression of reporters with CUC-CCG codon pairs (relative to reporter with all optimal codons).** However, complete removal of tL(UAG) is not predicted to eliminate CUC-CCG inhibition. Plot of GFP/RFP for one CUC-CCG inhibitory pair in strains with one, two or three copies of *tL(UAG)*.

**Supplemental Figure S5. Increased copies of tL(GAG) result in a linear increase in expression of reporters with two (IOIO) or four (IIII) CUC-CCG codon pairs (relative to reporter with all optimal codons).**

**Supplemental Figure S6.-Effects of *tP(UGG)* wild type and indicated mutants on relative expression from reporters with indicated inhibitory codon pairs at amino acids 5 or 6**. In each case, expression is compared to synonymous optimal reporter in same strain.

**Supplemental Figure S7.-Effects of *tP(UGG)* wild type and indicated mutants on relative expression from reporters with three copies of the indicated inhibitory codon pairs at amino acid 100 in *GLN4*(1-99)-(Codon Pair-NN)_3_ GFP reporter**. In each case, expression is compared to synonymous optimal reporter in same strain.

**Supplemental Figure S8. Mutations in *BCP1*, an essential chaperone for ribosomal protein Rpl23p suppress CUC-CCG inhibition.** (A) A suppressor strain with a mutation in *BCP1* restores growth on media lacking uracil and (B) exhibits increased GFP/RFP fluorescence from reporters with two CUC-CCG codon pairs.

**Supplemental Figure S9. Strains with reconstructed mutations in large ribosomal subunit proteins suppress inhibition by CUC-CCG.** GFP/RFP from indicated reporters in *tl(GAG)Δ* and *tL(GAG)Δ RPL7AΔ* and *tL(GAG)Δ RPL36BΔ* strains. (Data used in Fig. 4C).

**Supplemental Figure S10. Deletion of *RPL36B* suppresses inhibition by six inhibitory codon pairs but not by the three pairs responsive to *HEL2* and *MBF1***. GFP/RFP in wild type (blue) and *rpl36BΔ* (cyan) mutant from strains with optimal and inhibitory pairs. Synonymous pairs encoding the indicated dipeptide are grouped, with optimal pairs indicated in red and their synonymous inhibitory codon pairs in black. (Data used in Fig. 4D).

**Supplemental Figure S11. Loss of *RPL1B* strongly affects expression of a reporter with three copies of the CUC-CCG inhibitory codon pair**. GFP/RFP from indicated reporters in *tl(GAG)Δ* and *tL(GAG)Δ RPL1BΔ* strains. (Data used in Fig. 5A).

**Supplemental Figure S12. GFP from Leu-Pro variants with *PGK1* (short) and *YDL224C* (long) 5’ UTR. (blue *PGK1* UTR; pattern *YDL224C* UTR)**. GFP from reporters with four optimal codon pairs (OOOO: UUG-CCA), two inhibitory codon pairs (IOOI CUC-CCG;UUG-CCA) and three inhibitory codon pairs (IIOI,CUC-CCG;UUG-CCA). (Data used in Figure 5E.)

**Supplemental Figure S13. GFP from Arg-Arg variants with *PGK1* (short) and *YDL224C* (long) 5’ UTR. (blue *PGK1* UTR; pattern *YDL224C* UTR)**. GFP from reporters with four optimal codon pairs (OOOO: AGA-AGA), two inhibitory codon pairs (IOOI CGA-CGA, AGA-AGA) and three inhibitory codon pairs (IIOI, CGA-CGA, AGA-AGA). (Data used in Figure 5F.)

**Supplemental Figure S14. GFP/RFP expression of indicated reporters in wild type and reconstructed *SCH9* mutants.** Reconstructed *sch9 N543S* and *sch9 Q618\** mutants exhibit similar GFP/RFP from reporters with four optimal UUG-CCA codon pairs compared to a strain with wild type SCH9. However, the *sch9* mutants both exhibit substantially increased expression of a reporter with three inhibitory pairs compared to their parent strain with wild type *SCH9*. (Data used in Figure 6D).

**Supplemental Figure S15. Growth of *sch9* mutants obtained during selection of suppressors in *hel2Δ* or *mbf1Δ* parents.** Suppressor strains with mutations in *SCH9*, isolated as suppressors of *hel2Δ* and *mbf1Δ* parent strains, exhibit increased growth on media lacking uracil compared to their parents and reduced growth on YDP and YPG media.

**Supplemental Figure S16. GFP/RFP expression from CUC-CCG OIIO reporter in parents and mutant strains.** Suppressor strains with mutations in *SCH9*, isolated as suppressors of *hel2Δ* and *mbf1Δ* parent strains exhibit increased GFP/RFP expression from a reporter with two copies of the CUC-CCG inhibitory pair compared to their parents, although in some cases the increase is only slightly significant.

**Supplemental Figure S17.** Maps and sequences of EEVN391, EAH118-2 and EEVN535 vectors used for insertion of inhibitory pairs upstream of GFP (EEVN391), tRNAs and tRNA variants for insertion at LYS2 (EAH118-2), and insertion of inhibitory codon pairs upstream of URA3 (EEVN535) and EBB024, bearing a Flag tagged RFP-GFP fusion.

